# SARS-CoV-2 spike protein predicted to form complexes with host receptor protein orthologues from a broad range of mammals

**DOI:** 10.1101/2020.05.01.072371

**Authors:** SD Lam, N Bordin, VP Waman, HM Scholes, P Ashford, N Sen, L van Dorp, C Rauer, NL Dawson, CSM Pang, M Abbasian, I Sillitoe, SJL Edwards, F Fraternali, JG Lees, JM Santini, CA Orengo

**Affiliations:** Department of Applied Physics, Faculty of Science and Technology, Universiti Kebangsaan Malaysia, Bangi, Selangor, 43600, Malaysia; Institute of Structural and Molecular Biology, University College London, London, WC1E 6BT, UK; Indian Institute of Science Education and Research, Pune, 411008, India; UCL Genetics Institute, University College London, London, WC1E 6BT, UK; Department of Science and Technology Studies, University College London, London, WC1E 6BT, UK; Randall Division of Cell and Molecular Biophysics, Guy’s Campus, New Hunt’s House, King’s College London, London, SE1 1UL, UK; Department of Biological and Medical Sciences, Faculty of Health and Life Sciences, Oxford Brookes University, Oxford, OX3 OBP, UK

**Keywords:** SARS-CoV-2, COVID-19, spike protein, ACE2, TMPRSS2, structural bioinformatics

## Abstract

SARS-CoV-2 has a zoonotic origin and was transmitted to humans via an undetermined intermediate host, leading to infections in humans and other mammals. To enter host cells, the viral spike protein (S-protein) binds to its receptor, ACE2, and is then processed by TMPRSS2. Whilst receptor binding contributes to the viral host range, S-protein:ACE2 complexes from other animals have not been investigated widely. To predict infection risks, we modelled S-protein:ACE2 complexes from 215 vertebrate species, calculated changes in the energy of the complex caused by mutations in each species, relative to human ACE2, and correlated these changes with COVID-19 infection data. We also analysed structural interactions to better understand the key residues contributing to affinity. We predict that mutations are more detrimental in ACE2 than TMPRSS2. Finally, we demonstrate phylogenetically that human SARS-CoV-2 strains have been isolated in animals. Our results suggest that SARS-CoV-2 can infect a broad range of mammals, but few fish, birds or reptiles. Susceptible animals could serve as reservoirs of the virus, necessitating careful ongoing animal management and surveillance.

## Introduction

Severe acute respiratory syndrome coronavirus 2 (SARS-CoV-2) is a novel coronavirus that emerged towards the end of 2019 and is responsible for the coronavirus disease 2019 (COVID-19) global pandemic. Available data suggests that SARS-CoV-2 has a zoonotic source[1], with the closest sequence currently available deriving from the horseshoe bat[2]. As yet, the transmission route to humans, including the intermediate host, is unknown. So far, little work has been done to assess the animal reservoirs of SARS-CoV-2, or the potential for the virus to spread to other species living with, or in close proximity to, humans in domestic, rural, agricultural or zoological settings.

Coronaviruses, including SARS-CoV-2, are major multi-host pathogens and can infect a wide range of non-human animals[3–5]. SARS-CoV-2 is a member of the Betacoronavirus genus, which includes viruses that infect economically important livestock, including cows[6] and pigs[7], together with mice[8], rats[9], rabbits[10], and wildlife, such as antelope and giraffe[11]. Severe acute respiratory syndrome coronavirus (SARS-CoV), the betacoronavirus that caused the 2002-2004 SARS outbreak[12], likely jumped to humans from its original bat host via civets. Viruses genetically similar to human SARS-CoV have been isolated from animals as diverse as racoon dogs, ferret-badgers[3] and pigs[13], suggesting the existence of a large host reservoir. It is therefore probable that SARS-CoV-2 can also infect a wide range of species.

Real-world SARS-CoV-2 infections have been reported in cats[14], lions and tigers[15], dogs[16,17] and minks[16,17]. Animal infection studies have also identified cats[18] and dogs[18] as hosts, as well as ferrets[18], macaques[19] and marmosets[19]. Recent *in vitro* studies have also suggested an even broader set of animals may be infected[20–22]. To understand the potential host range of SARS-CoV-2, the plausible extent of zoonotic and anthroponotic transmission, and to guide surveillance efforts, it is vital to know which species are susceptible to SARS-CoV-2 infection.

The receptor binding domain (RBD) of the SARS-CoV-2 spike protein (S-protein) binds to the extracellular peptidase domain of angiotensin I converting enzyme 2 (ACE2) mediating cell entry[23]. The sequence of ACE2 is highly conserved across vertebrates, suggesting that SARS-CoV-2 could use orthologues of ACE2 for cell entry. The structure of the SARS-CoV-2 S-protein RBD has been solved in complex with human ACE2[24]. Identification of critical binding residues in this structure have provided valuable insights into viral recognition of the host receptor[24–29]. Deep mutagenesis studies have also revealed residues important for stability[30,31]. Compared with SARS-CoV, the SARS-CoV-2 S-protein has a 10–22-fold higher affinity for human ACE2[24,25,32], due to more contacts in the interface that cover a larger surface area[29], and three mutational hotspots in the S-protein that lead to a more specific and compact conformation [29,33]. Similarly, variations in human ACE2 have also been found to increase affinity for S-protein receptor binding [34]. These factors may contribute to the host range and infectivity of SARS-CoV-2.

Both SARS-CoV-2 and SARS-CoV additionally require the transmembrane serine protease (TMPRSS2) to mediate cell entry. Together, ACE2 and TMPRSS2 confer specificity of host cell types that the virus can enter[35,36]. Upon binding to ACE2, the S-protein is cleaved by TMPRSS2 at two cleavage sites on separate loops, which primes the S-protein for cell entry[37]. TMPRSS2 has been docked against the SARS-CoV-2 S-protein, which revealed its binding site to be adjacent to these two cleavage sites[36]. An approved TMPRSS2 protease inhibitor drug is able to block SARS-CoV-2 cell entry[38], which demonstrates the key role of TMPRSS2 alongside ACE2[39]. As such, both ACE2 and TMPRSS2 represent attractive therapeutic targets against SARS-CoV-2[40].

Recent work has predicted possible hosts for SARS-CoV-2 using the structural interplay between the S-protein and ACE2. These studies proposed a broad range of hosts, covering hundreds of mammalian species, including tens of bat[27] and primate[41] species, and more comprehensive studies analysing all classes of vertebrates[41–43], including agricultural species of cow, sheep, goat, bison and water buffalo. In addition, sites in ACE2 have been identified as under positive selection in bats, particularly in regions involved in binding the S-protein[27,44]. The impacts of mutations in ACE2 orthologues have also been tested, for example structural modelling of ACE2 from 27 primate species[41] demonstrated that apes and African and Asian monkeys may also be susceptible to SARS-CoV-2. However, whilst cell entry is necessary for viral infection, it may not be sufficient alone to cause disease. For example, variations in other proteins may prevent downstream events that are required for viral replication in a new host. Hence, examples of real-world infections[14–17] and experimental data from animal infection studies[18–22] are required to validate hosts that are predicted to be susceptible.

Here, we analysed the effect of known mutations in orthologues of ACE2 and TMPRSS2 from a broad range of 215 vertebrate species, including primates, rodents and other placental mammals; birds; reptiles; and fish. For each species, we generated a 3-dimensional model of the ACE2 protein structure from its protein sequence and calculated the impacts of known mutations in ACE2 on the stability of the S-protein:ACE2 complex. We correlated changes in the energy of the complex with changes in the structure of ACE2, chemical properties of residues in the binding interface, and experimental COVID-19 infection phenotypes from *in vivo* and *in vitro* animal studies. To further test our predictions and rationalise the key sites contributing to energy changes of the complex, we performed detailed manual structural analyses, presented as a variety of case studies for different species. Unlike other studies that analyse interactions that the S-protein makes with the host, we also analyse the impact of mutations in vertebrate orthologues of TMPRSS2. Our results suggest that SARS-CoV-2 could infect a broad range of vertebrates, which could serve as reservoirs of the virus, supporting future anthroponotic and zoonotic transmission.

## Results

### Conservation of ACE2 in vertebrates

We aligned protein sequences of 247 vertebrate orthologues of ACE2. Most orthologues have more than 60% sequence identity with human ACE2 (Supplementary Fig. 1A). For each orthologue, we generated a 3-dimensional model of the protein structure from its protein sequence using FunMod[45,46]. We were able to build high-quality models for 236 vertebrate orthologues, with nDOPE scores < −1 (Supplementary Table 5). 11 low-quality models were removed from the analysis.

**Figure 1:**
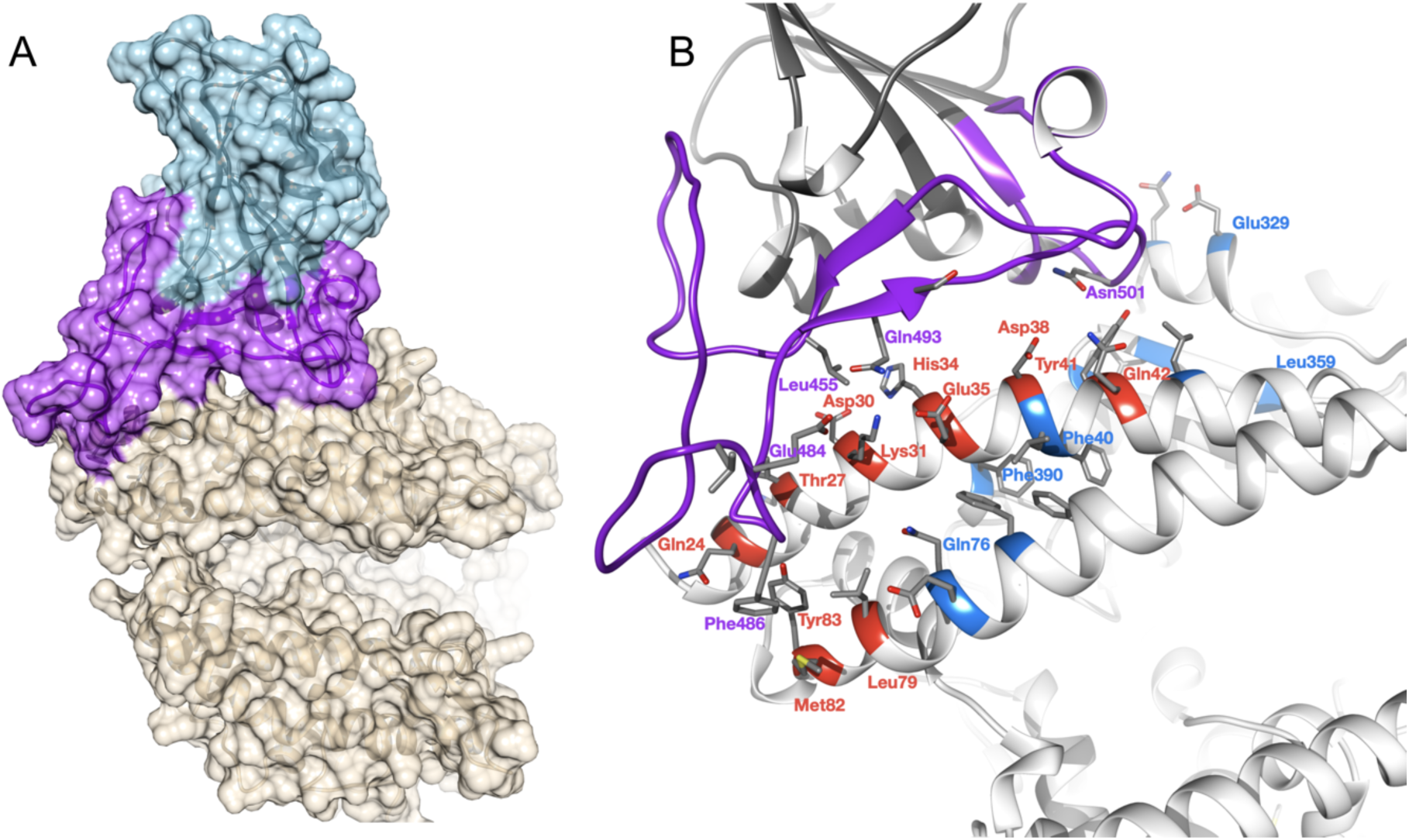
Overview of SARS-CoV-2 S-protein:human ACE2 complex and interface. (a) SARS-CoV-2 S-protein RBD (light blue, purple) showing the receptor binding motif (purple) at the interface with ACE2 (tan). (b) residues in the SARS-CoV-2 S-protein:human ACE2 complex interface. Key RBD interface residues are shown (purple) with a subset of ACE2 contact residues that are not conserved in vertebrates for DC (red) and DCEX residues (blue) (PDB ID 6M0J).

**Figure 2:**
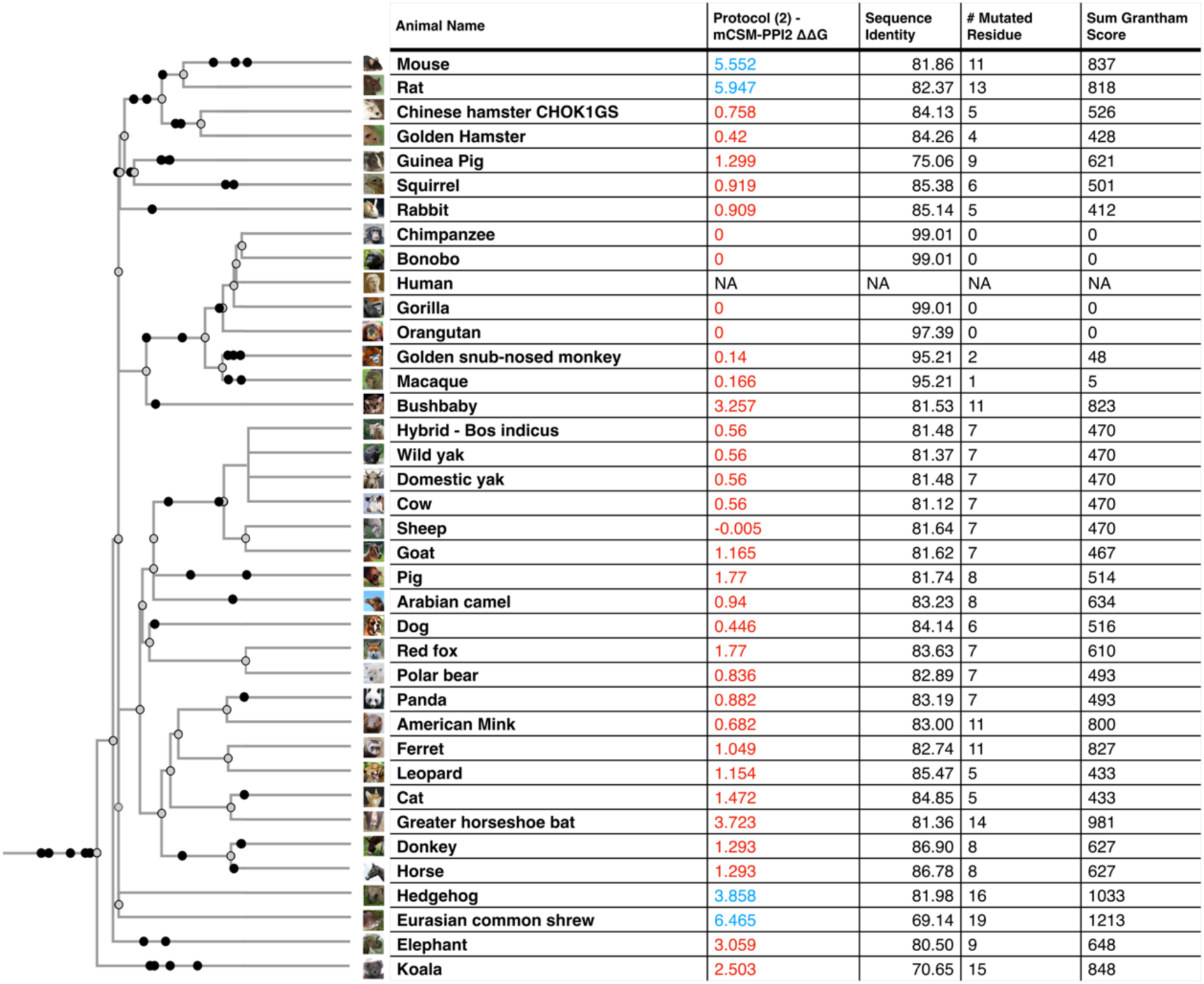
Phylogenetic tree of species that humans come into close contact with in domestic, agricultural or zoological settings. Leaves are annotated by the change in energy of the complex (ΔΔG), as measured by protocol 2 mCSM-PPI2. Animals are categorised according to risk of infection by SARS-CoV-2, with ΔΔG ≤ 3.7 being at risk (red), and ΔΔG > 3.7 not at risk (blue). These thresholds were chosen as they agree well with the available experimental data (Fig. 3). For each animal, the number of residue changes compared to human ACE2 sequence and the total chemical shift (measured by Grantham score - see Supplementary Methods 5) across the DCEX residues are also shown. This tree contains a subset of animals from Supplementary Fig. 7. Animal photos courtesy of ENSEMBL and associated sources (https://www.ensembl.org/info/about/image_credits.html)

### Identification of critical S-protein:ACE2 interface residues

ACE2 residues directly contacting the S-protein (DC residues) were identified in a structure of the complex (PDB ID 6M0J; Fig. 1a, Supplementary Results 2, Supplementary Fig. 3). We also identified a more extended set of both DC residues and residues within 8Å of DC residues likely to be influencing binding (DCEX residues).

**Figure 3:**
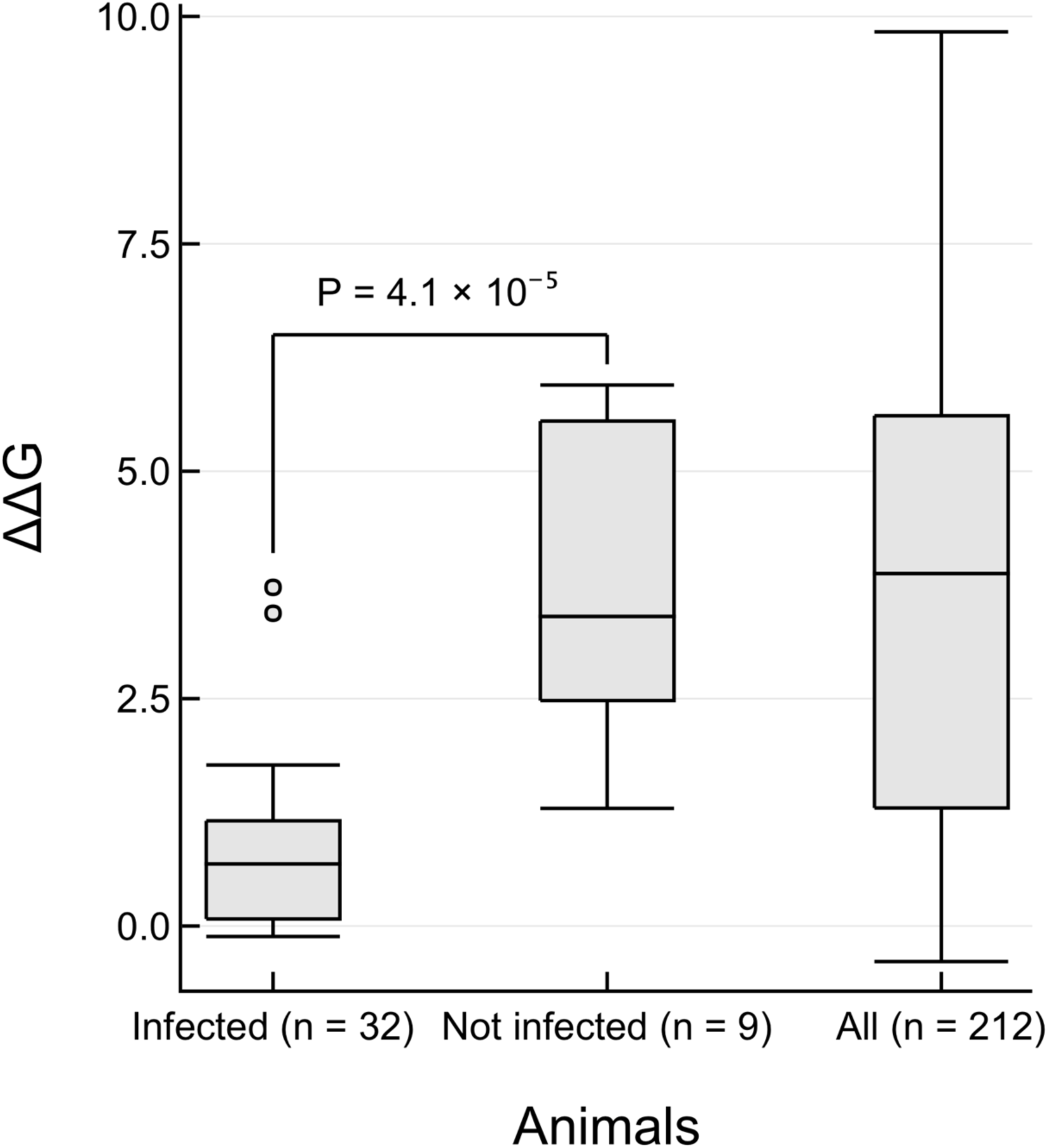
Changes in energy of S-protein:ACE2 complex for animals that can be infected by SARS-CoV-2. Boxplots of ΔΔG values calculated by protocol 2 mCSM-PPI2 are shown. *Infected:* 32 animals that have *in vivo* or *in vitro* or real world evidence of infection. *Not infected:* 9 animals that have been experimentally tested but show no infection. *All:* 212 animals that were included in this study. The one-sided *P* value is reported from a Mann-Whitney test of the hypothesis that ΔΔG values from infected animals is lower than for not infected animals, against the null hypothesis that there is no difference between the two distributions.

After analysing the orthologue interfaces, we removed models that were missing > 10 DCEX residues. 21 models were removed from the analysis, leaving 215 models to take forward for further analysis. We observed high sequence (> 60% identity) and structure similarity (score > 90 out of 100) between ACE2 proteins for all species (Supplementary Results 1).

### Changes in the energy of the S-protein:ACE2 complex in vertebrates

We used multiple methods to assess the relative change in binding energy (ΔΔG) of the SARS-CoV-2 S-protein:ACE2 complex following mutations in DC residues and DCEX residues that are likely to influence binding. We found that protocol 2 employing mCSM-PPI2 (henceforth referred to as P(2)-PPI2), calculated over the DCEX residues, correlated best with the phenotype data (Supplementary Results 3, Supplementary Fig. 4, Table 1), justifying the use of animal models to calculate ΔΔG values in this context. Since this protocol considers mutations from animal to human, lower ΔΔG values correspond to stabilisation of the animal complex relative to the human complex, and therefore higher risk of infection. We show the residues that P(2)-PPI2 reports as stabilising or destabilising for the SARS-CoV-2 S-protein:ACE2 animal complex for DC (Supplementary Fig. 8) and DCEX (Supplementary Fig. 9) residues. To consider ΔΔG values in an evolutionary context, we annotated phylogenetic trees for all 215 vertebrate species analysed (Supplementary Fig. 7) and for a subset of animals that humans come into close contact with in domestic, agricultural or zoological settings (Fig. 2).

**Table 1:** Collated evidence of in vivo, in vitro and real world animal infections to date [14–17,19,21,22,48–51]. ΔΔG values calculated by protocol 2 (mCSM-PPI2) and Grantham scores are also shown. Cell colours denote animals that have been infected (red), not infected (blue) or no experimental evidence (grey). Animals are categorised according to risk of infection by SARS-CoV-2, with ΔΔG ≤ 3.72 being at risk (red), and ΔΔG > 3.72 not at risk (blue). These thresholds were chosen as they agree well with the available experimental data (Fig. 3).

| Animal | Evidence of infection | In vivo infection | In vitro infection | Real world infection | $\Delta\Delta G$ | Grantham score |
| --- | --- | --- | --- | --- | --- | --- |
| Baboon | 1 |  | 1<sup>48</sup> |  | -0.115 | 5 |
| Bat (horseshoe) | 1 |  | 1<sup>21 22 48</sup> |  | 3.723 | 981 |
| Bear | 1 |  | 1<sup>48</sup> |  | 0.044 | 493 |
| Buffalo | 1 |  | 1<sup>48</sup> |  |  |  |
| Camel | 1 |  | 1<sup>21</sup> |  | 0.940 | 634 |
| Capuchin | 0 |  | 0<sup>48</sup> |  | 3.404 | 280 |
| Cat | 1 | 1<sup>49</sup> | 1<sup>21 22 48</sup> | 1<sup>14</sup> | 1.472 | 433 |
| Chicken | 0 | 0<sup>49</sup> | 0<sup>21</sup> |  | 5.001 | 1350 |
| Chimp | 1 |  | 1<sup>48</sup> |  | 0.000 | 0 |
| Civet | 1 |  | 1<sup>21 22</sup> |  |  |  |
| Colobus | 1 |  | 1<sup>48</sup> |  | 0.000 | 0 |
| Cow | 1 |  | 1<sup>21 48</sup> |  | 0.560 | 470 |
| Dog | 1 | 1<sup>49</sup> | 1<sup>22 48</sup> | 1<sup>16 17</sup> | 0.446 | 516 |
| Dolphin | 1 |  | 1<sup>48</sup> |  | 1.399 | 548 |
| Donkey | 0 |  | 0<sup>21</sup> |  | 1.293 | 627 |
| Duck | 0 | 0<sup>49</sup> |  |  | 5.889 | 1394 |
| Ferret | 1 | 1<sup>49 50</sup> |  |  | 1.049 | 827 |
| Fox | 1 |  | 1<sup>48</sup> |  | 1.770 | 610 |
| Gelada | 1 |  | 1<sup>48</sup> |  | -0.055 | 5 |
| Gibbon | 1 |  | 1<sup>48</sup> |  | 0.089 | 26 |
| Goat | 1 |  | 1<sup>21 48</sup> |  | 1.165 | 467 |
| Golden snub-nosed monkey | 1 |  | 1<sup>48</sup> |  | 0.140 | 48 |
| Gorilla | 1 |  | 1<sup>48</sup> |  | 0.000 | 0 |
| Guinea pig | 0 |  | 0<sup>21</sup> |  | 1.299 | 621 |
| Hamster | 1 |  | 1<sup>48</sup> |  | 0.420 | 526 |
| Horse | 1 |  | 1<sup>21 48</sup> |  | 1.293 | 627 |
| Jerboa | 1 |  | 1<sup>48</sup> |  |  |  |
| Koala | 0 |  | 0<sup>48</sup> |  | 2.503 | 848 |
| Leopard | 1 |  | 1<sup>48</sup> |  | 1.154 | 433 |
| Lynx | 1 |  | 1<sup>48</sup> |  | 0.734 | 433 |
| Macaques | 1 | 1<sup>19 51</sup> | 1<sup>48</sup> |  | 0.166 | 5 |
| Marmoset | 1 | 1<sup>19</sup> | 0<sup>48</sup> |  | 3.438 | 280 |
| Mink | 1 |  |  | 1<sup>16 17</sup> | 0.632 | 800 |
| Mouse | 0 |  | 1<sup>21 22 48</sup> |  | 5.552 | 837 |
| White footed mouse | 1 |  | 1<sup>48</sup> |  |  |  |
| Orangutan | 1 |  | 1<sup>48</sup> |  | 0.000 | 0 |
| Panda | 1 |  | 1<sup>48</sup> |  | 0.882 | 493 |
| Pangolin | 1 |  | 1<sup>21 22 48</sup> |  |  |  |
| Pig | 1 | 0<sup>49</sup> | 1<sup>21 48</sup> |  | 1.770 | 514 |
| Puma | 1 |  | 1<sup>48</sup> |  |  |  |
| Rabbit | 1 |  | 1<sup>21 22 48</sup> |  | 0.909 | 412 |
| Rat | 0 |  | 0<sup>21 22</sup> |  | 5.947 | 818 |
| Rhinoceros | 1 |  | 1<sup>48</sup> |  |  |  |
| Roussette | 1 |  | 1<sup>48</sup> |  |  |  |
| Hawaiian monk seal | 0 |  | 0<sup>48</sup> |  |  |  |
| Sealion | 1 |  | 1<sup>48</sup> |  |  |  |
| Sheep | 1 |  | 1<sup>21 48</sup> |  | -0.055 | 470 |
| Stoat | 0 |  | 0<sup>48</sup> |  |  |  |
| Squirrel | 1 |  | 1<sup>48</sup> |  | 0.919 | 501 |
| Squirrel monkey | 0 |  | 0<sup>48</sup> |  | 2.479 | 280 |
| Tiger | 1 |  |  | 1<sup>15</sup> |  |  |
| Whales | 1 |  | 1<sup>48</sup> |  | 0.784 | 563 |
| Yak | 1 |  | 1<sup>48</sup> |  | 0.560 | 470 |

**Figure 4:**
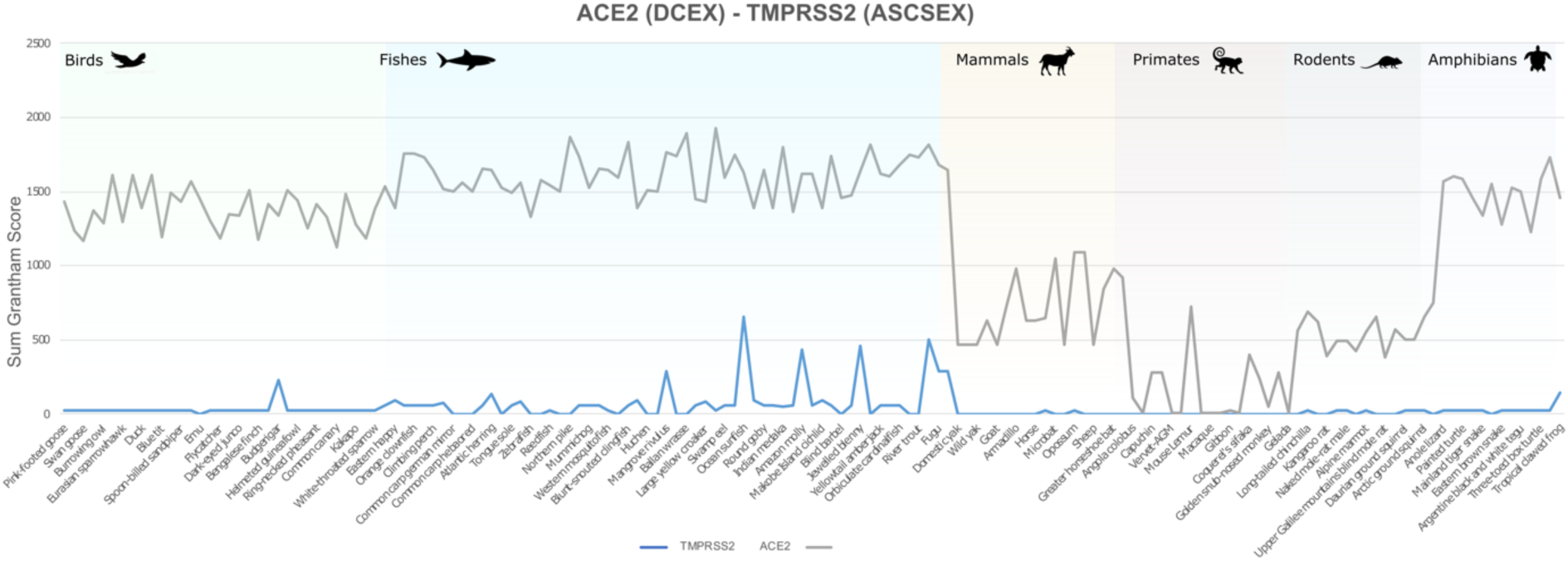
Comparison of Grantham score sums for ASCSEX residues in ACE2 and TMPRSS2.

In general we see a high infection risk for most mammals, with a notable exception for all non-placental mammals. ΔΔG values measured by P(2)-PPI2 correlate well with the infection phenotypes (Table 1). ΔΔG values are significantly lower for animals that can be infected by SARS-CoV-2 than for animals for which there is no evidence of infection (Fig. 3; Mann-Whitney one-sided *P* = 4.1 × 10^−5^). Two animals are outliers in the infected boxplot, corresponding to horseshoe bat (ΔΔG = 3.723) and marmoset (ΔΔG = 3.438). To be cautious, since *in vivo* experiments have shown that marmosets can be infected, and *in vitro* experiments have shown that horseshoe bats can be infected[2,20–22] (Table 1), we consider animals that have ΔΔG values less than, or equal to, the ΔΔG = 3.7 for horseshoe bat to be at risk. Additionally, there is a clear sampling bias in the set of animals that have so far been experimentally characterised: all but chicken and duck are mammals. As more non-mammals are tested, the median ΔΔG value for non-infection is likely to increase. In further support of these predictions we analysed the 41 animals having experimental evidence using an orthogonal method, HADDOCK [47], and found ∼95% agreement between the two independent approaches for animals predicted to be at risk (see Supplementary Results 3 and Supplementary Figure 6).

**Figure 5:**
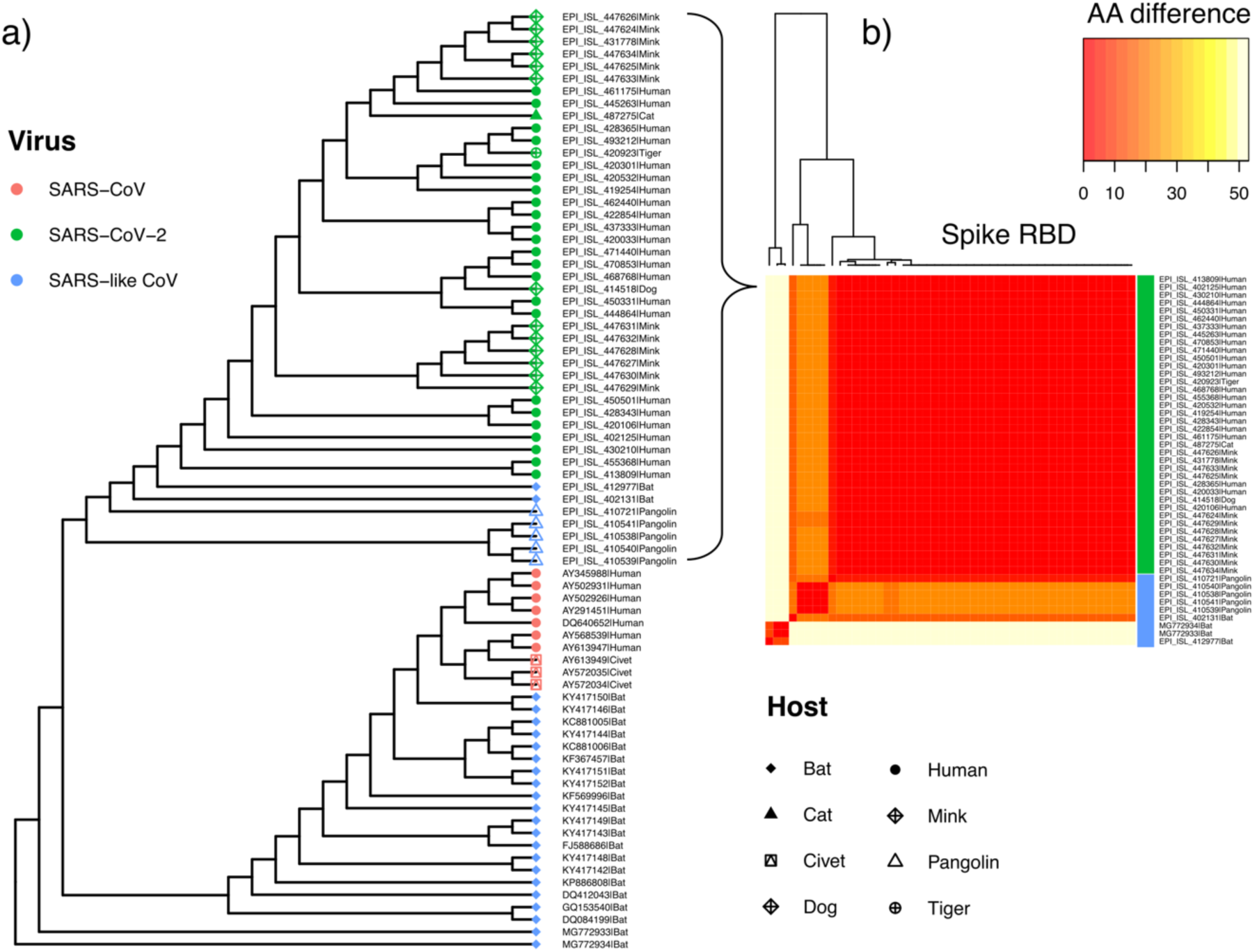
Phylogeny of SARS-like viruses. (a) Genome-wide maximum likelihood phylogenetic tree of SARS-like betacoronavirus strains sampled from diverse hosts (coloured tip symbols provide host and species; Supplementary Table 3). Genomes EPI_ISL_402131 and EPI_ISL_412977 are samples from RaTG13 and RmYN02 isolated from horseshoe bat hosts (Rhinolophus affinis and R malayanus respectively). (b) Pairwise amino acid differences (color scale) at the S-protein RBD between human and animal associated strains of SARS-CoV-2, relative to closely related SARS-like viruses in bat and pangolin hosts.

**Figure 6:**
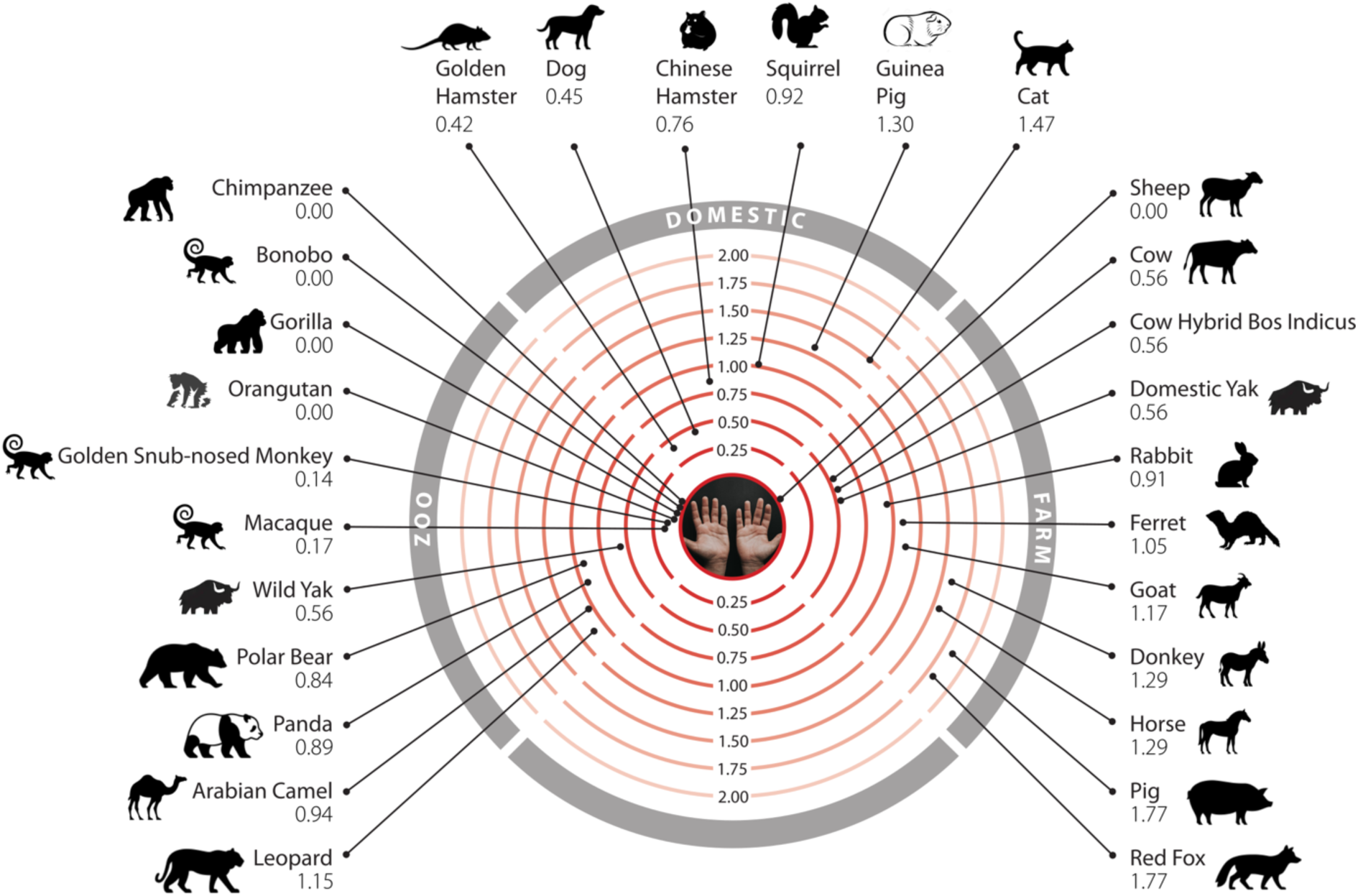
Mammals that humans come into contact with that are at risk of infection by SARS-CoV-2. Twenty-six mammals are categorised into domestic, *agricultural or zoological settings.* Numbers represent the change in binding energy (ΔΔG) of the S-protein:ACE2.

As shown in previous studies, and supported by experimental data, many primates are predicted to be at high risk[19,41,42]. In agricultural settings, camels, cows, sheep, goats and horses also have relatively low ΔΔG values, suggesting comparable binding affinities to humans, in agreement with experimental data[20,21]. In domestic settings, dogs[18], cats[18], hamsters[20], and rabbits[20–22] also have ΔΔG values suggesting risk, again in agreement with experimental data (Table 1). Whilst, zoological animals that come into contact with humans, such as pandas, leopards and bears, are also at risk of infection as shown experimentally[20] and suggested by our calculated ΔΔG values. Importantly, mice and rats do not appear to be susceptible (ΔΔG values > 3.7), so hamsters and ferrets are being used as model organisms for human COVID-19. Of the 35 birds tested only a handful, including the blue tit, show an infection risk. Similarly, out of 72 fish in this study, relatively few show a low change in energy of the complex, suggesting that most have no susceptibility to infection. Those susceptible include the common carp, turbot and Nile tilapia. Of the 14 reptiles and amphibians investigated, only turtle and crocodile show any risk.

In predicting susceptibility, we have chosen thresholds supported by *in vivo* or *in vitro* experimental data. Previous work contrasted the binding energy of the S-protein of SARS-CoV and SARS-CoV-2 with human ACE2 protein[24,25,32]. SARS-CoV is able to infect humans despite a ∼20-fold lower binding affinity[24,25,32], suggesting that even where mutations in different animal species make the interfaces less compatible for SARS-CoV2, a considerably decreased binding energy may still be sufficient to enable infection. By applying this threshold we correctly predict all 32 animals in our dataset that have experimental evidence of infection, to be at risk (Table 1).

However, for a few animals we predict at risk using this threshold, *in vitro* experimental studies to date have not shown infection. For example, donkeys are at risk of infection (ΔΔG = 1.3) but no infections were observed *in vitro* for these animals[21]. However, infection has been observed *in vitro* for horse[21] and horse and donkey have identical DCEX residues and the same ΔΔG. Amongst New World monkeys, marmosets have been experimentally infected[19]. We predict that the closely related capuchin and squirrel monkey are also at risk, although they have not been shown to be infected using functional assays[20]. We performed detailed structural analyses to characterise the key residues contributing to binding energy changes and to consider these discrepancies further. Our analyses reveal that the interfaces in both capuchin and squirrel monkey are similar to marmoset, suggesting that these two New World monkeys are also likely to be at risk even though there is no current experimental data supporting this[20] (Supplementary Results 4). Furthermore, all these monkeys have high global sequence similarity to human. For capuchin and squirrel monkey this is >∼90% and their DCEX residues are identical to those of human, further supporting risk. In marmoset, which has experimental evidence of infection the global sequence identity is 89% and 93% over the DCEX residues.

Additionally, we compared changes in energy of the S-protein:ACE2 complex in SARS-CoV-2 and SARS-CoV and found similar changes suggesting that the range of animals susceptible to the virus is likely to be similar for SARS-CoV-2 and SARS-CoV (Supplementary Results 5).

### Conservation of TMPRSS2 and its role in SARS-CoV-2 infection

ACE2 and TMPRSS2 are key factors in the SARS-CoV-2 infection process. Both are highly co-expressed in susceptible cell types, such as type II pneumocytes in the lungs, ileal absorptive enterocytes in the gut, and nasal goblet secretory cells[52]. Since both proteins are required for infection of host cells, and since our analyses clearly support suggestions of conserved binding of S-protein:ACE2 across animal species, we decided to analyse whether the TMPRSS2 was similarly conserved. There is no known structure of TMPRSS2, so we built a high-quality model (nDOPE = −0.78) from a template structure (PDB ID 5I25). Since TMPRSS2 is a serine protease, and the key catalytic residues are known, we used FunFams[53] to identify highly conserved residues in the active site and the cleavage site that are likely to be involved in substrate binding. This resulted in two sets of residues that we analysed: the active site and cleavage site residues (ASCS), and the active site and cleavage site residues plus residues within 8Å of catalytic residues that are highly conserved in the FunFam (ASCSEX). The sum of Grantham scores for mutations in the active site and cleavage site for TMPRSS2 is zero or consistently lower than ACE2 in all organisms under consideration, for both ASCS and ASCSEX residues (Fig. 4). This means that the mutations in TMPRSS2 involve more conservative changes.

Mutations in DCEX residues seem to have a more disruptive effect in ACE2 than in TMPRSS2. Whilst we expect orthologues from organisms that are close to humans to be conserved and have lower Grantham scores, we observed some residue substitutions that have high Grantham scores for primates, such as capuchin, marmoset and mouse lemur. In addition, primates, such as the coquerel sifaka, greater bamboo lemur and Bolivian squirrel monkey, have mutations in DCEX residues with high Grantham scores. Mutations in TMPRSS2 may render these animals less susceptible to infection by SARS-CoV-2.

### Phylogenetic Analysis of SARS-like strains in different animal species

A small-scale phylogenetic analysis was performed on a subset of SARS-CoV-2 assemblies in conjunction with a broader range of SARS-like betacoronaviruses (Supplementary Table 4), including SARS-CoV isolated from both humans and civets. Consistent with previous phylogenetic work[2], SARS-like viruses isolated from horseshoe bats (RaTG13, EPI_ISL_402131; RmYN02, EPI_ISL_412977) are the closest relatives of SARS-CoV-2 strains currently available in genomic repositories (Fig. 5), though still remain many decades divergent from SARS-CoV-2[54]. Aided by a large community sequencing effort, tens of thousands of human-associated SARS-CoV-2 genome assemblies are now accessible on GISAID[16,17]. At the time of writing, these also include one complete assembly generated from a virus infecting a domestic dog (EPI_ISL_414518), one genome obtained from a zoo tiger (EPI_ISL_420923), one directly isolated from a domestic cat (EPI_ISL_487275) and 12 high coverage complete genomes obtained from farmed mink (including EPI_ISL_431778). SARS-CoV-2 strains from animal infections fall among the phylogenetic diversity observed in a representative set of human strains (Fig. 5a), as also seen in larger phylogenetic analyses available on NextStrain (https://nextstrain.org/ncov/global). Irrespective of host, the SARS-CoV-2 spike receptor binding domain is conserved (Fig. 4b) across tested human and animal associated SARS-CoV-2, suggesting mutations in the RBD are not required for infections observed in non-human species to date. Of note, whilst genome-wide data indicates a closer phylogenetic relationship between SARS-CoV-2 strains and species in circulation in horseshoe bats, the receptor binding domain alignment instead supports a closer relationship with a SARS-like virus isolated from pangolins[55] (EPI_ISL_410721; Fig. 5b), in line with previous reports[56].

## Discussion

The ongoing COVID-19 global pandemic has a zoonotic origin, necessitating investigations into how SARS-CoV-2 infects animals, and how the virus can be transmitted across species. Given the role that the stability of the complex, formed between the S-protein and its receptors, could contribute to the viral host range, zoonosis and anthroponosis, there is a clear need to study these interactions. However, to our knowledge there have been few studies of relative changes in the energies of the S-protein:ACE2 complex [43]. A number of recent studies[20,21,42] have suggested that, due to high conservation of ACE2, some animals are vulnerable to infection by SARS-CoV-2. Concerningly, these animals could, in theory, serve as reservoirs of the virus, increasing the risk of future zoonotic events, though transmission rates across species are currently not known. Therefore, it is important to try to predict which other animals could potentially be infected by SARS-CoV-2, so that the plausible extent of transmission can be estimated, and surveillance efforts can be guided appropriately.

Animal susceptibility to infection by SARS-CoV-2 has been studied *in vivo*[18,19,50,51,57] and *in vitro*[20–22] during the course of the pandemic. Parallel *in silico* work has made use of the protein structure of the S-protein:ACE2 complex to computationally predict the breadth of possible viral hosts. Most studies simply considered the number of residues mutated relative to human A[32,48,58]57], although some also analyse the effect that these mutations have on the interface stabil[34,41,59]58]. The most comprehensive of these studies analysed the number and locations of mutated residues in ACE2 orthologues from 410 species[42], but did not perform detailed energy calculations as we have done. Few studies have explored changes in the energy of the S-protein:ACE2 complex on a large scale. Shortly after we reported our work in BioRxiv Rodrigues et al. [43] submitted a paper in BioRxiv also reporting changes in binding energy of the complex for 30 different animal species, measured using a different approach (HADDOCK [47]). The results are in good agreement with ours (nearly 90% of the risk assessments agree). Furthermore, when we applied HADDOCK to the 41 animals for which experimental data exists, we also observed significant agreement in risk assessments with those predicted using mCSM-PPI2 (see Supplementary Results 3, Supplementary Figure 6 and Supplementary Methods 4). Our HADDOCK analysis showed slightly better correlation with experiment than the Rodrigues et al. study, possibly due to use of a different template structure when building the animal models (6M0J, a better resolved structure than 6M17 used by Rodrigues et al.). Our work is the only study that has so far explored changes in the energy of the S-protein:ACE2 complex on a very large scale (215 animals) in order to assess risk of infection across a broad range of animal species. Furthermore, it is the only study to assess whether changes in TMPRSS2 could also be influencing risk.

In this study, we performed a comprehensive analysis of the major proteins that SARS-CoV-2 uses for cell entry. We predicted structures of ACE2 and TMPRSS2 orthologues from 215 vertebrate species and modelled S-protein:ACE2 complexes. We calculated relative changes in energy (ΔΔG) of S-protein:ACE2 complexes, *in silico*, following mutations from animal residues to those in human. Our predictions suggest that, whilst many mammals are susceptible to infection by SARS-CoV-2, most birds, fish and reptiles are not likely to be. However, there are some exceptions. We manually analysed residues in the S-protein:ACE2 interface, including DC residues that directly contacted the other protein, and DCEX residues that also included residues within 8Å of the binding residues, that may affect binding. We clearly showed the advantage of performing more sophisticated studies of the changes in energy of the complex, over more simple measures––such as the number or chemical nature of mutated residues––used in other studies. Furthermore, the wider set of DCEX residues that we identified near the binding interface had a higher correlation to the phenotype data than the DC residues. In addition to ACE2, we also analysed how mutations in TMPRSS2 impact binding to the S-protein. We found that mutations in TMPRSS2 are less disruptive than mutations in ACE2, indicating that binding interactions in the S-protein:TMPRSS2 complex in different species will not be affected.

To increase our confidence in assessing changes in the energy of the complex, we developed multiple protocols using different, established methods. We correlated these stability measures with experimental infection phenotypes in the literature, from *in vivo*[18,19,50,51] and *in vitro*[20–22] studies of animals. Protocol 2 using mCSM-PPI2 (P(2)-PPI2) correlated best with the number of mutations, chemical changes induced by mutations and infection phenotypes, so we chose to focus our analysis employing this protocol. Our method cannot determine relative changes in energy that are associated with no risk. Instead, we used experimental *in vivo* and *in vitro* infection data as the gold standard to identify animals at risk. Of note, horseshoe bats, heavily advocated as a putative reservoir host, are predicted to be infected from *in vitro* experiments, despite the considerable disruption in the interface that our detailed structural analysis shows.

We found that our predicted ΔΔG values for animals that can be infected by SARS-CoV-2 are significantly lower than for animals that showed no infection when tested experimentally (Fig. 1, Table 1). ΔΔG values for horseshoe bat and marmoset were outliers for infected animals. These ΔΔG values are higher than the median ΔΔG value for animals that are not infected and are approximately the same value as the median ΔΔG = 3.88 for all animals included in this study. However, this may be a result of the biased sampling of animals that have been tested experimentally, where most have been mammals to date. Going forward, if more distantly related animals are experimentally characterised, it is plausible that non-placental animals, of which many have ΔΔG > 3.7 (the value obtained for horseshoe bat), would be found to not be infected. Therefore, the difference between ΔΔG values for animals that can, and cannot, be infected by SARS-CoV-2 will increase. Overall, our measurements of the change in energy of the complex for the SARS-CoV S-protein were highly correlated with SARS-CoV-2, so our findings are also applicable to SARS-CoV.

Humans are likely to come into contact with 26 of these species in domestic, agricultural or zoological settings (Fig. 6). Of particular concern are sheep, that have no change in energy of the S-protein:ACE2 complex, as these animals are farmed and come into close contact with humans. Indeed, SARS-CoV-2 is already responsible for infections in various animal species. SARS-CoV-2 genomes[16,17] have been isolated from natural infections in zoo lions and tigers[15], companion animals including cats and dogs[60,61] and following widespread outbreaks in multiple mink farms in the Netherlands resulting in mass culling[62] (Fig. 5). In most cases natural infections have been linked to human infections supporting cross-species transmission and high levels of exposure[14]. To date, minks provide the only well supported example of sustained intraspecies transmission with secondary zoonotic transmission back to humans[62]. Consistently, we predict American mink to be at risk of infection by SARS-CoV-2, with ΔΔG = 0.632.

To gain a better understanding of the nature of the S-protein:ACE2 interface, we performed more detailed structural analyses for a subset of species. In a few cases, we had found discrepancies between our energy calculations and experimental phenotypes, namely predicting risk for some animals where *in vitro* experiments showed no infection (Table 1). To test our predictions, we manually analysed how the shape or chemistry of residues may impact complex stability for all DC residues and a selection of DCEX residues. Previous studies have identified a number of important locations in human ACE2 for binding the S-protein[25,29] and we found agreement with these in structural studies using our animal models. These locations, namely the hydrophobic cluster near the N-terminus and two hotspot locations near residues 31 and 353, stabilise the binding interface. SARS-CoV-2 exploits the hydrophobic pocket by mutations that alter the conformation and flexibility of the RBD loop, together with point mutation L486F that provides a compact interface which is more dynamically and energetically favourable compared with SARS-CoV[25,29]. Our structural analysis showed how SARS-CoV-2 can utilise this pocket for binding at the interface in all the species we examined, including those for which current experimental test data suggest no risk.

Hotspot 353 shows more structural variability. In agreement with our calculations of large changes in energy of the S-protein:ACE2 complex in horseshoe bat, our structural studies show that the variant D38N causes the loss of a salt bridge and H-bonding interactions between ACE2 and S-protein at hotspot 353. These detailed structural analyses are supported by the high Grantham score and calculated total ΔΔG for the change in energy of the complex. Both dog and cat have a physico-chemically similar variant at this hotspot (D38E), which although disrupting the salt bridge still permits alternative H-bonding interactions between the spike RBD and ACE2. For marmosets and other New World monkeys, capuchin and squirrel monkey, our structural analyses revealed similarity to human at hotspot 353 in the ACE2 interface. In fact, capuchin resembled the human ACE2 interface even more closely than marmoset, which can be infected[19], even though *in vitro* experiments have not reported infection in capuchin[20]. Our ΔΔG value for capuchin (ΔΔG = 3.4) suggests risk of infection, and also for squirrel monkey (ΔΔG = 2.5), despite the fact that squirrel monkey also failed to show risk in *in vitro* experimental studies. Of note is the fact that marmoset showed no infection in *in vitro* studies[20], whilst recent *in vivo* experiments[19] have shown risk, perhaps suggesting that it can be difficult to detect infection *in vitro* for these monkeys. Alternatively, the lack of infection may suggest additional factors influencing infection and indicate that these animals, which are primates closely related to human, may be useful models for studying immune, or other factors, related to resistance.

Finally, our structural analyses showed that some DCEX residues were likely to be allosteric sites, which may represent promising drug targets[63].

The value of our study is not in determining an absolute ΔΔG threshold for risk, but rather in providing information about relative changes in binding energy that will allow the host range of the virus to be more accurately gauged once more experimental work has been conducted. We believe that false positive predictions are more acceptable than false negatives. So, within the context of possible transmission events between species, and particularly to human, we consider that an animal can be infected if there is any experimental evidence of infection.

We applied protocols that enabled a comprehensive study of host range, within a reasonable time, for identifying species at risk of infection by SARS-CoV-2, or of becoming reservoirs of the virus. Although we felt that these faster methods were justified by the need for timely answers to these questions, there are clearly caveats to our work that should be taken into account. Whilst we use a state of the art modelling tool[64] and an endorsed method for calculating changes in energy of the complex[65], molecular dynamics may give a more accurate picture of energy changes by sampling rotamer space more comprehensively[29]. However, such an approach would have been prohibitively expensive at a time when it is clearly important to identify animals at risk as quickly as possible. Each animal could take orders of magnitude longer to analyse using molecular dynamics. Further caveats include the fact that although the animals we highlight at risk from our changes in binding energy calculations correlate well with the experimental data, there is only a small amount of such data currently available, and many of the experimental papers reporting these data are yet to be peer reviewed. Finally, we restricted our analyses to one strain of SARS-CoV-2, but other strains may have evolved with mutations that give more complementary interfaces. For example, recent work suggests SARS-CoV-2 can readily adapt to infect mice following serial passages[66].

In summary, our work is not aiming to provide an absolute measure of risk of infection. Rather, it should be considered an efficient method to screen a large number of animals and suggest possible susceptibility, and thereby guide further studies. Any predictions of possible risk should be confirmed by experimental studies and computationally expensive, but more robust methods, like molecular dynamics.

The ability of SARS-CoV-2 to infect host cells and cause COVID-19, sometimes resulting in severe disease, ultimately depends on a multitude of other host-virus protein interactions[40]. While we do not investigate them all in this study, our results suggest that SARS-CoV-2 could indeed infect a broad range of mammals. As there is a possibility of creating new reservoirs of the virus, we should now consider how to identify such transmission early and to mitigate against such risks. In particular, farm animals and other animals living in close contact with humans could be monitored, protected where possible and managed accordingly[67].

## Methods

### Sequence Data

ACE2 protein sequences for 239 vertebrates, including humans, were obtained from ENSEMBL[68] version 99 and eight sequences from UniProt release 2020_1 (Supplementary Table 1). TMPRSS2 protein sequences for 278 vertebrate sequences, including the human sequence, were obtained from ENSEMBL (Supplementary Table 2).

A phylogenetic tree of species, to indicate the evolutionary relationships between animals, was downloaded from ENSEMBL[68].

### Structural Data

The structure[24] of the SARS-CoV-2 S-protein bound to human ACE2 at 2.45Å was used throughout (PDB ID 6M0J).

### Sequence analysis

We used standard methods to analyse the sequence similarity between human ACE2 and other vertebrate species (Supplementary Methods 1). We also mapped ACE2 and TMPRSS2 sequences to our CATH functional families to detect residues highly conserved across species (Supplementary Methods 1).

### Structure analysis

#### Identifying residues in ACE2

In addition to residues in ACE2 that contact the S-protein directly, various other studies have also considered residues that are in the second shell, or are buried, and could influence binding[30]. Therefore, in our analyses we built on these approaches and extended them to compile the following sets for our study:

1. *Direct contact (DC)* residues. This includes a total of 20 residues that are involved in direct contact with the S-protein[24] identified by PDBe[69] and PDBSum[70].
2. *Direct Contact Extended (DCEX) residues*. This dataset includes residues within 8Å of DC residues, that are likely to be important for binding. These were selected by detailed manual inspection of the complex, and also considering the following criteria: (i) reported evidence from deep mutagenesis[30], (ii) *in silico* alanine scanning (using mCSM-PPI[71]), (iii) residues with high evolutionary conservation patterns identified by the FunFam-based protocol described above, i.e. residues identified with DOPS ≥ 70 and ScoreCons score ≥ 0.7, (iv) allosteric site prediction (Supplementary Methods 2), and (v) sites under positive selection (Supplementary Methods 2). Selected residues are shown in Supplementary Fig. 3 and residues very close to DC residues (i.e. within 5Å) are annotated.

We also included residues identified by other related structural analyses, reported in the literature (Supplementary Methods 2).

### Generating 3-dimensional structure models

Using the ACE2 protein sequence from each species, structural models were generated for the S-protein:ACE2 complex for 247 animals using the FunMod modelling pipeline[45,46] (Supplementary Methods 3). FunMod searches for structural templates by mapping sequences to a CATH FunFam and selecting the structure of the closest relative of known structure, to use as a template for homology modelling [53]. Sequences are mapped by scanning them against the CATH FunFam HMM library using HMMer3 [72]. The structural template selected was PDB ID 6M0J, a high-resolution crystal structure of SARS-CoV2 S-protein:human ACE2 complex. We generated query–template alignments using HH-suite[73] and predicted 3D models using MODELLER v.9.24 [64]. The ‘very_slow’ schedule was used for model refinement to optimise the geometry of the complex and interface. For each species, we generated 10 models and selected the model with the lowest nDOPE[74] score. Only high-quality models were used in this analysis, with nDOPE score < −1 and with < 10 DCEX residues missing. This gave a final dataset of 215 animals for further analysis.

The modelled structures of ACE2 were compared against the human structure (PDB ID 6M0J) and pairwise, against each other, using SSAP[75]. SSAP measures the similarity between 3D protein structures by calculating similarity in vector views between aligned residues. A vector view for a given residue is the set of vectors from the Cβ atom of that residue to the Cβ atom of all other residues in the protein structure. SSAP returns a score in the range 0-100, with identical structures scoring 100 [73].

We also built models for TMPRSS2 proteins in all available species and identified the residues likely to be involved in the protein function (see Supplementary Methods 3).

### Measuring changes in the energy of the S-protein:ACE2 complex in SARS-CoV-2 and SARS-CoV

We calculated the changes in binding energy of the SARS-CoV-2 S-protein:ACE2 complex and the SARS-CoV S-protein:ACE2 complex of different species, compared to human, following two different protocols:

1. *Protocol 1:* Using the human complex and mutating the residues for the ACE2 interface to those found in the given animal sequence and then calculating the ΔΔG of the complex using both mCSM-PPI1[71] and mCSM-PPI2[65] (Supplementary Methods 4). This gave a measure of the destabilisation of the complex in the given ***animal relative to the human*** complex. ΔΔG values < 0 are associated with destabilising mutations, whilst values ≥ 0 are associated with stabilising mutations.
2. *Protocol 2:* We repeated the analysis with both mCSM-PPI1 and mCSM-PPI2 as in protocol 1, but using the animal 3-dimensional models, instead of the human ACE2 structure, and calculating the ΔΔG of the complex by mutating the animal ACE2 interface residue to the appropriate residue in the human ACE2 structure. This gave a measure of the destabilisation of the complex in the ***human complex relative to the given animal***. Values ≤ 0 are associated with destabilisation of the human complex (i.e. animal complexes more stable), whilst values > 0 are associated with stabilisation of the human complex (i.e. animal complexes less stable).

We subsequently correlated ΔΔG values with available *in vivo* and *in vitro* experimental data on COVID-19 infection data for mammals. Protocol 2, mCSM-PPI2, correlated best with these data.

### Change in residue chemistry for mutations

To measure the degree of chemical change associated with mutations occurring in DC and DCEX residues, we computed the Grantham score[76] for each vertebrate compared to the human sequence (Supplementary Methods 5).

### Phylogeny of SARS-like betacoronaviruses

We performed phylogenetic analyses for a subset of SARS-CoV (n = 10), SARS-like (n = 28) and SARS-CoV-2 (n = 38) viruses from publicly available data in NCBI[77–84] and GISAID[16,17] (Supplementary Methods 6).

## Data Availability

The FunMod structural models for the SARS-CoV-2 Spike-RBD:ACE2 complex and TMPRSS are available on Zenodo at https://zenodo.org/record/3963980 [85].

## Supporting information

Supplentary Table S4 - SARS-CoV-2 Genomes metadata

Supplementary Table S1 - ACE2 Orthologs

Supplementary Table S2 - TMPRSS2 Orthologs

Supplementary Table S3 - Grantham scores averages and sums for ACE2 and TMPRSS2 in vertebrates

Supplementary Table S5 - nDOPE scores

## Acknowledgments

We thank Gal Horesh, Caitlin Lee Carpenter and Mohd Firdaus Raih for insightful discussions; Alan Hunns for help in making figures; and Laurel Woodridge, Sean Le Cornu and Declan Torin Cook for comments on the manuscript. We would also like to thank Francois Balloux, whose team member, Lucy van Dorp, contributed the phylogenetic analysis of SARS-like viruses.

## Author contributions

SL conceived the idea of analysing structures and effects of mutations in the S-protein:human ACE2 complex. JL conceived the idea of extending the analyses to animal complexes, for animals reported to be infected. JS conceived the idea of extending to a larger set of animals to explore host range. CO conceived the idea of contrasting multiple protocols to validate predictions. SL, CO, JL, JS designed the experiments. SL, NB, VW, PA, LvD performed the experiments. SL, NB, VW, HS, PA, NS, JS, LvD, CR, IS, JL, CO analysed data. SL, NB, VW, HS, PA, NS, JS, LvD, CR, ND, IS, JL, CO interpreted the results. SL, NB, VW, HS, PA, NS, JS, LvD, CR, ND, CSMP, MA, IS, JL, CO contributed to the manuscript and figures. HS, VW, PA, SL, CO wrote the manuscript. JL, SE, FF, JS, CO revised the manuscript.

