## Supplementary Figure S7 for "SARS-CoV-2 spike protein predicted to form complexes with host receptor protein orthologues from a broad range of mammals"

***Animal photos courtesy of ENSEMBL and associated sources (https://www.ensembl.org/info/about/image_credits.html)***

### Primates

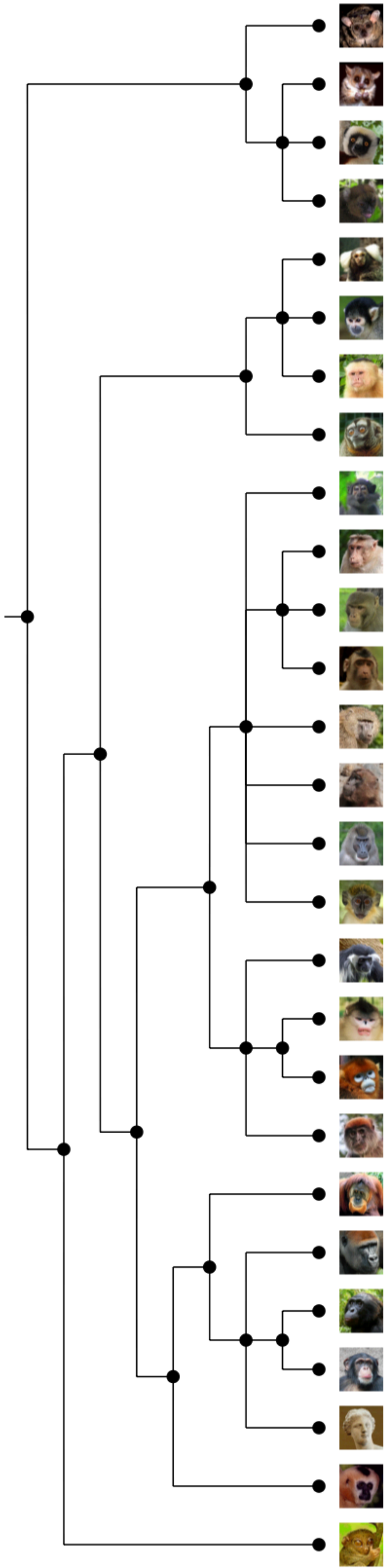

| Animal Name | Protocol (2) - mCSM-PPI2 ΔΔG | Sequence Identity | # Mutated Residue | Sum Grantham Score |
|---|---|---|---|---|
| Bushbaby | 3.257 | 81.53 | 11 | 823 |
| Mouse Lemur | 2.821 | 82.22 | 8 | 727 |
| Coquerel's sifaka | 0.28 | 85.86 | 4 | 404 |
| Greater bamboo lemur | 1.127 | 84.85 | 4 | 239 |
| **Marmoset** | 3.438 | 87.44 | 4 | 280 |
| Bolivian squirrel monkey | 2.479 | 92.05 | 4 | 280 |
| **Capuchin** | 3.404 | 92.17 | 4 | 280 |
| Ma's night monkey | 2.932 | 92.17 | 4 | 280 |
| Sooty mangabey | -0.243 | 95.21 | 1 | 5 |
| **Crab-eating macaque** | 0.166 | 95.21 | 1 | 5 |
| **Macaque** | 0.166 | 95.21 | 1 | 5 |
| **Pig-tailed macaque** | -0.267 | 95.34 | 1 | 5 |
| **Olive baboon** | -0.155 | 95.34 | 1 | 5 |
| **Gelada** | -0.055 | 95.34 | 1 | 5 |
| Drill | -0.297 | 94.96 | 1 | 5 |
| Vervet-AGM | 0.153 | 94.84 | 1 | 5 |
| **Angola colobus** | 0.013 | 95.88 | 2 | 107 |
| Black snub-nosed monkey | NIL | NIL | NIL | NIL |
| **Golden snub-nosed monkey** | 0.14 | 95.21 | 2 | 48 |
| **Ugandan red Colobus** | 0 | 95.03 | 0 | 0 |
| **Orangutan** | 0 | 97.39 | 0 | 0 |
| **Gorilla** | 0 | 99.01 | 0 | 0 |
| Bonobo | 0 | 99.01 | 0 | 0 |
| **Chimpanzee** | 0 | 99.01 | 0 | 0 |
| **Human** | NA | NA | NA | NA |
| **Gibbon** | 0.089 | 97.64 | 1 | 26 |
| Tarsier | 3.159 | 84.10 | 9 | 561 |

### Rabbit and Rodents (1)

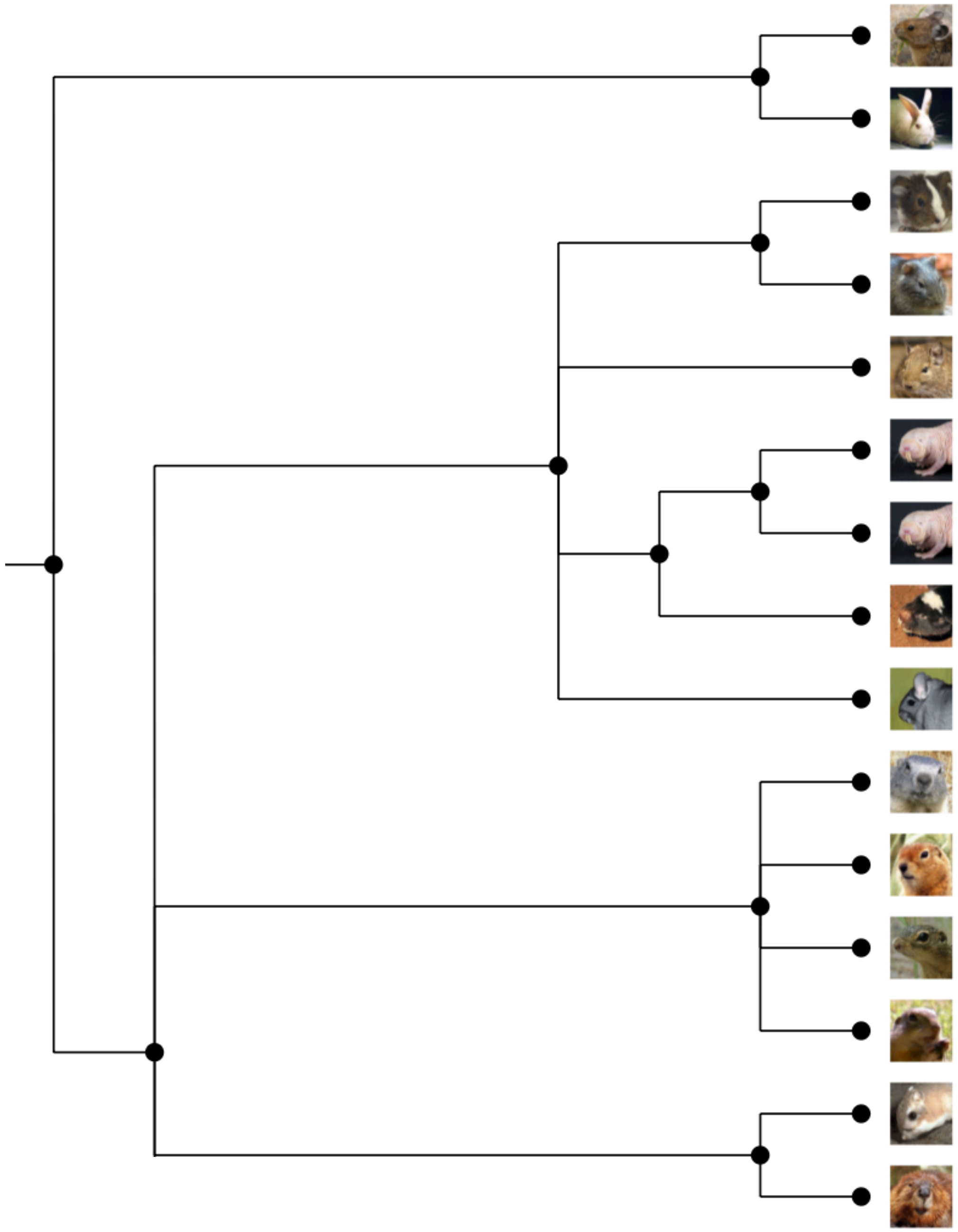

| Animal Name | Protocol (2) - mCSM-PPI2 ΔΔG | Sequence Identity | # Mutated Residue | Sum Grantham Score |
|---|---|---|---|---|
| Pika | NIL | NIL | NIL | NIL |
| **Rabbit** | 0.909 | 85.14 | 5 | 412 |
| **Guinea Pig** | 1.299 | 75.06 | 9 | 621 |
| Brazilian Guinea Pig | NIL | NIL | NIL | NIL |
| Degu | 1.953 | 84.47 | 8 | 629 |
| Naked mole-rat male | 1.065 | 84.47 | 6 | 493 |
| Naked mole-rat female | 1.065 | 84.60 | 6 | 493 |
| Damara mole rat | 1.84 | 84.72 | 7 | 578 |
| Long-tailed chinchilla | 3.339 | 84.72 | 9 | 688 |
| Alpine marmot | 1.833 | 84.88 | 7 | 557 |
| Arctic ground squirrel | 2.258 | 84.76 | 7 | 660 |
| **Squirrel** | 0.919 | 85.38 | 6 | 501 |
| **Daurian ground squirrel** | 0.876 | 84.88 | 6 | 501 |
| Kangaroo rat | -0.391 | 82.12 | 5 | 390 |
| American beaver | NIL | NIL | NIL | NIL |

### Rodents (2)

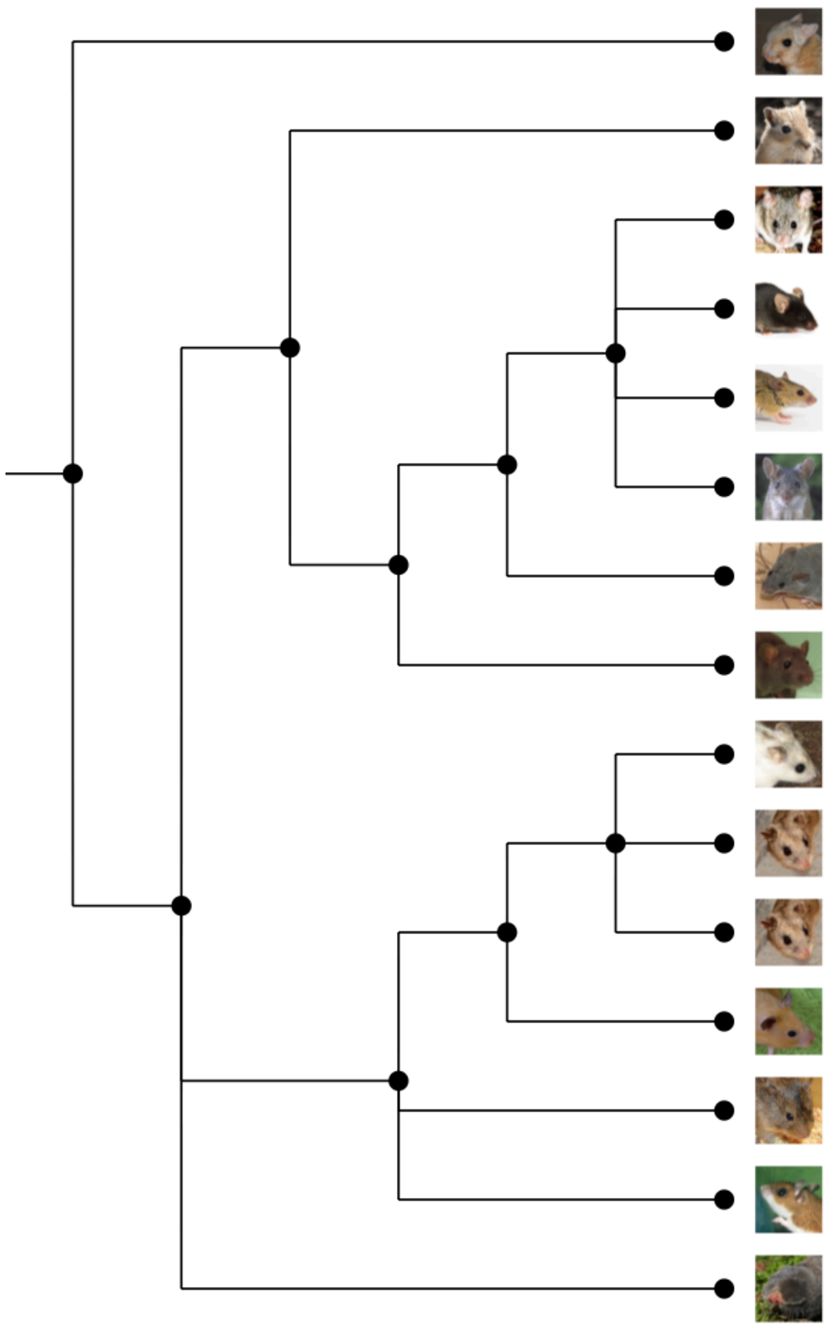

| Animal Name | Protocol (2) - mCSM-PPI2 ΔΔG | Sequence Identity | # Mutated Residue | Sum Grantham Score |
|---|---|---|---|---|
| **Lesser Egyptian jerboa** | 1.321 | 84.64 | 7 | 524 |
| Mongolian gerbil | NIL | NIL | NIL | NIL |
| Ryukyu mouse | 5.264 | 81.99 | 10 | 743 |
| Mouse | 5.552 | 81.86 | 11 | 837 |
| Algerian mouse | 3.653 | 82.37 | 10 | 785 |
| Steppe mouse | 4.556 | 78.34 | 11 | 841 |
| Shrew mouse | 4.611 | 82.87 | 10 | 750 |
| Rat | 5.947 | 82.37 | 13 | 818 |
| **Chinese hamster CHOK1GS** | 0.758 | 84.13 | 5 | 526 |
| **Chinese hamster CRIGRI** | 0.758 | 84.13 | 5 | 526 |
| **Chinese hamster PICR** | 0.758 | 84.01 | 5 | 526 |
| **Golden Hamster** | 0.42 | 84.26 | 4 | 428 |
| Prairie vole | 2.582 | 83.63 | 8 | 656 |
| Northern American deer mouse | 2.435 | 83.38 | 6 | 571 |
| Upper Galilee mountains blind mole rat | 0.281 | 82.37 | 4 | 381 |

### Mammals (1)

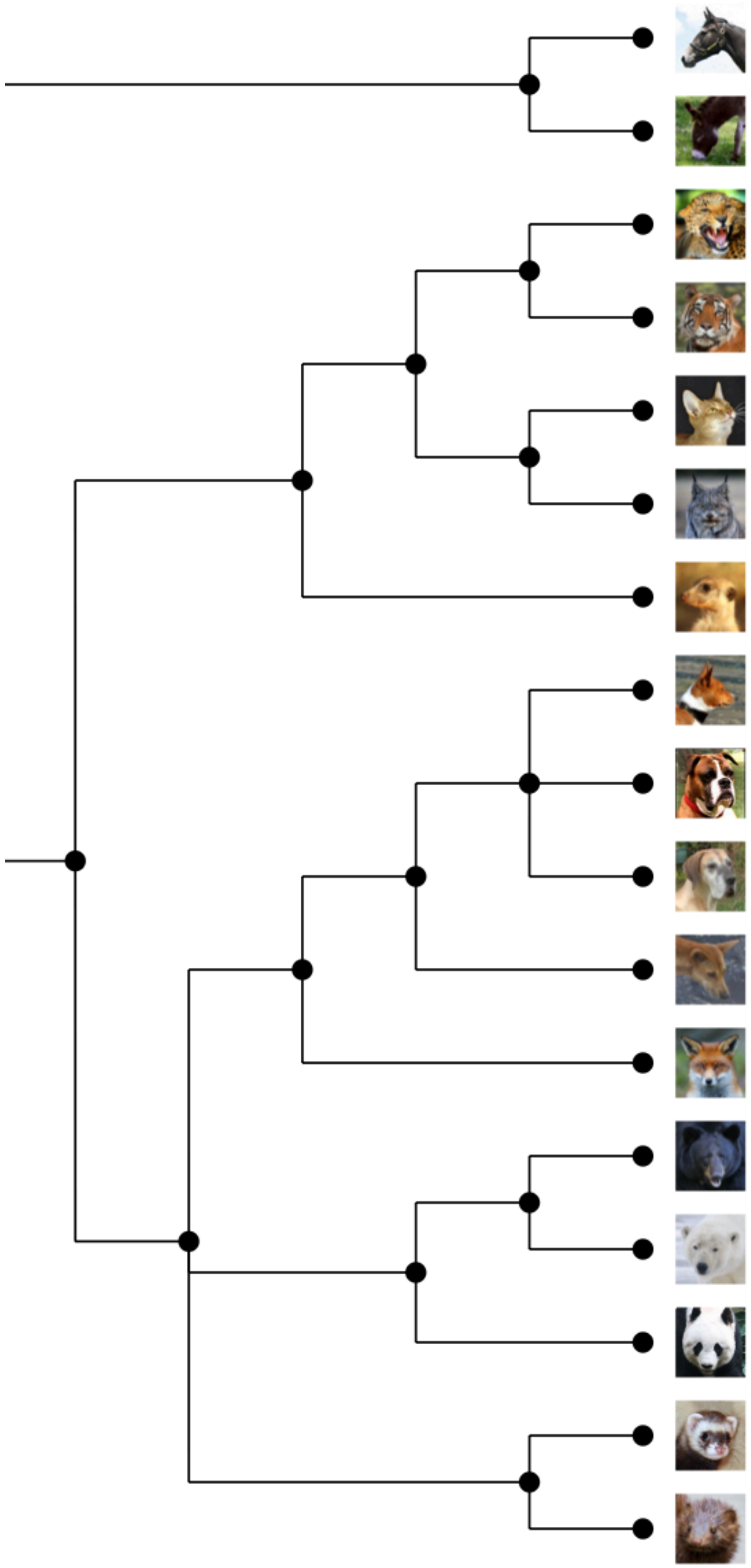

| Animal Name | Protocol (2) - mCSM-PPI2 ΔΔG | Sequence Identity | # Mutated Residue | Sum Grantham Score |
|---|---|---|---|---|
| **Horse** | 1.293 | 86.78 | 8 | 627 |
| Donkey | 1.293 | 86.90 | 8 | 627 |
| **Leopard** | 1.154 | 85.47 | 5 | 433 |
| **Tiger** | NIL | NIL | NIL | NIL |
| **Cat** | 1.472 | 84.85 | 5 | 433 |
| **Canada lynx** | 0.734 | 85.09 | 5 | 433 |
| Meerkat | 1.963 | 82.73 | 12 | 921 |
| Dog - Basenji | NIL | NIL | NIL | NIL |
| **Dog** | 0.446 | 84.14 | 6 | 516 |
| Dog - Great Dane | 0.446 | 84.14 | 6 | 516 |
| Dingo | -0.136 | 84.01 | 6 | 516 |
| **Red fox** | 1.77 | 83.63 | 7 | 610 |
| **American black bear** | 0.044 | 84.01 | 7 | 493 |
| **Polar bear** | 0.836 | 82.89 | 7 | 493 |
| **Panda** | 0.882 | 83.19 | 7 | 493 |
| **Ferret** | 1.049 | 82.74 | 11 | 827 |
| **American mink** | 0.632 | 83.00 | 11 | 800 |

### Mammals (2)

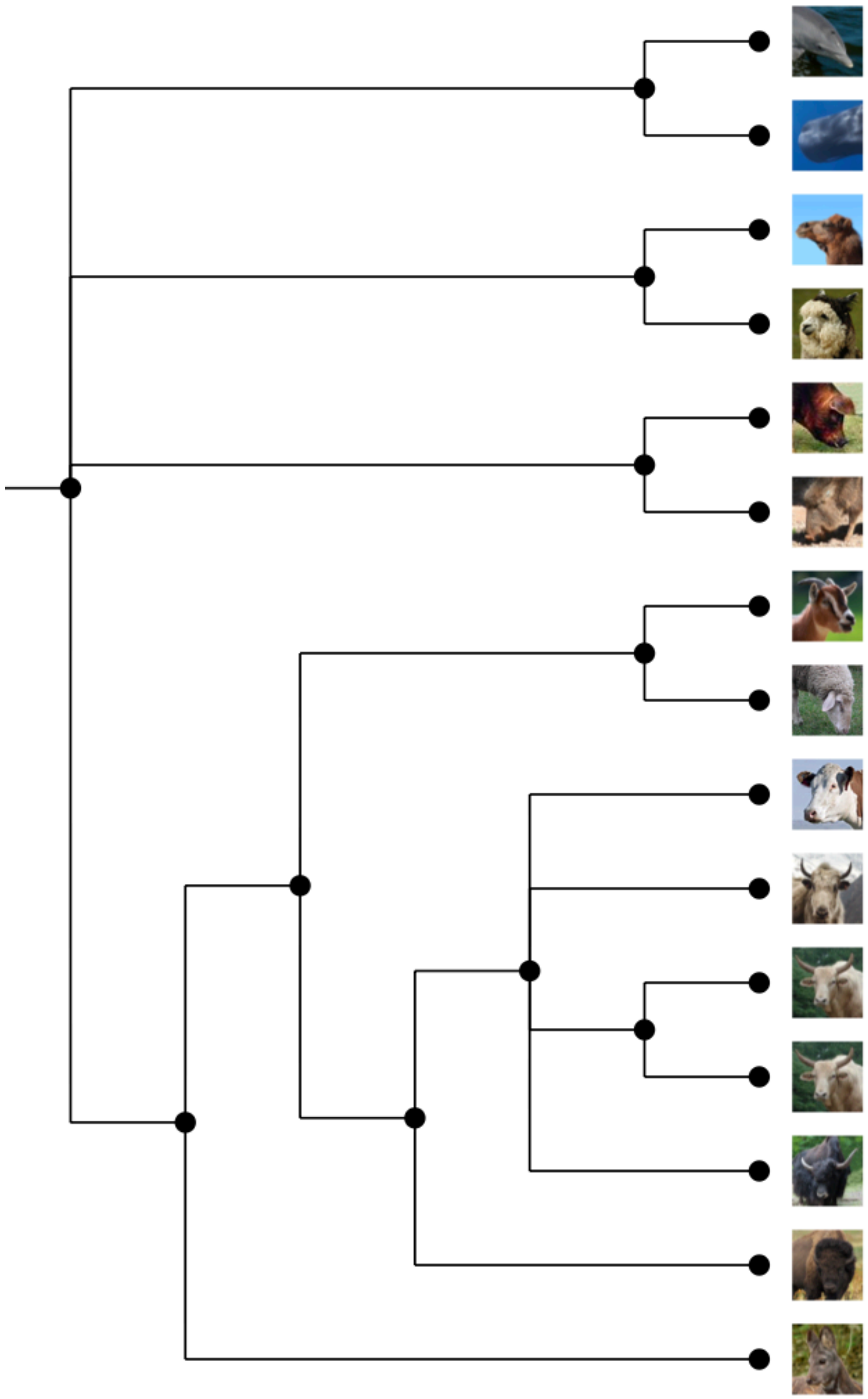

| Animal Name | Protocol (2) - mCSM-PPI2 ΔΔG | Sequence Identity | # Mutated Residue | Sum Grantham Score |
|---|---|---|---|---|
| **Dolphin** | 1.399 | 75.65 | 9 | 548 |
| **Sperm whale** | 0.784 | 82.73 | 7 | 563 |
| **Arabian camel** | 0.94 | 83.23 | 8 | 634 |
| Alpaca | NIL | NIL | NIL | NIL |
| **Pig** | 1.77 | 81.74 | 8 | 514 |
| Chacoan peccary | 1.416 | 81.99 | 10 | 720 |
| **Goat** | 1.165 | 81.62 | 7 | 467 |
| **Sheep** | -0.005 | 81.64 | 7 | 470 |
| **Cow** | 0.56 | 81.12 | 7 | 470 |
| **Domestic yak** | 0.56 | 81.48 | 7 | 470 |
| Hybrid - Bos Taurus | NIL | NIL | NIL | NIL |
| **Hybrid - Bos Indicus** | 0.56 | 81.48 | 7 | 470 |
| **Wild yak** | 0.56 | 81.37 | 7 | 470 |
| American bison | NIL | NIL | NIL | NIL |
| Siberian musk deer | -0.349 | 81.24 | 7 | 470 |

### Mammals (3)

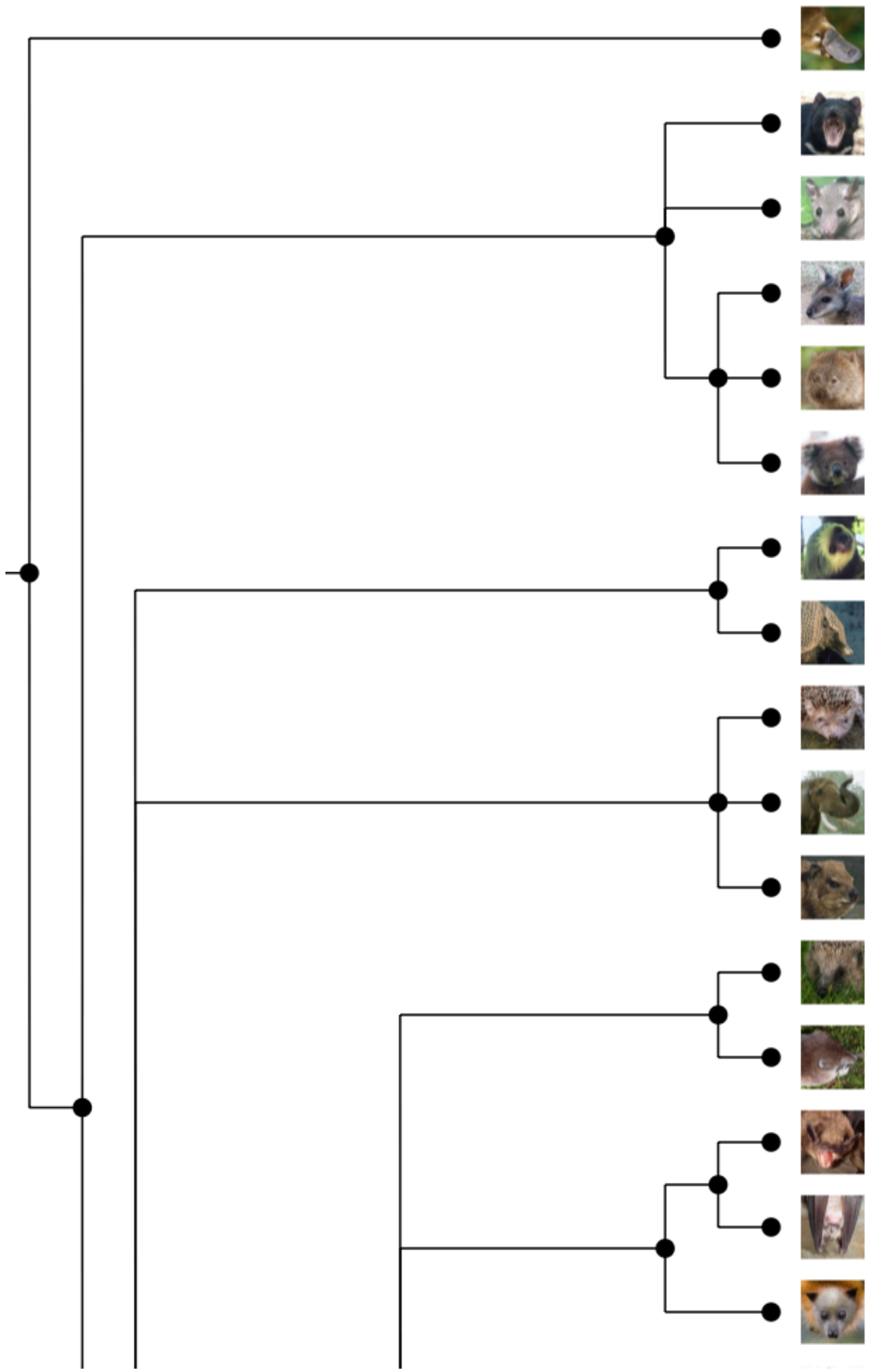

| Animal Name | Protocol (2) - mCSM-PPI2 ΔΔG | Sequence Identity | # Mutated Residue | Sum Grantham Score |
|---|---|---|---|---|
| Platypus | 4.083 | 68.14 | 17 | 1092 |
| Tasmanian devil | NIL | NIL | NIL | NIL |
| Opossum | 5.332 | 71.20 | 16 | 1094 |
| Wallaby | NIL | NIL | NIL | NIL |
| Common wombat | 2.751 | 71.68 | 16 | 870 |
| Koala | 2.503 | 70.65 | 15 | 848 |
| Sloth | NIL | NIL | NIL | NIL |
| Armadillo | 5.364 | 78.71 | 16 | 981 |
| Lesser hedhehog tenrec | NIL | NIL | NIL | NIL |
| Elephant | 3.059 | 80.50 | 9 | 648 |
| Hyrax | NIL | NIL | NIL | NIL |
| Hedgehog | 3.858 | 81.98 | 16 | 1033 |
| Eurasian common shrew | 6.465 | 69.14 | 19 | 1213 |
| Microbat | 4.429 | 80.45 | 15 | 1050 |
| **Greater horseshoe bat** | 3.723 | 81.36 | 14 | 981 |
| Megabat | 1.962 | 79.75 | 9 | 374 |

### Birds (1)

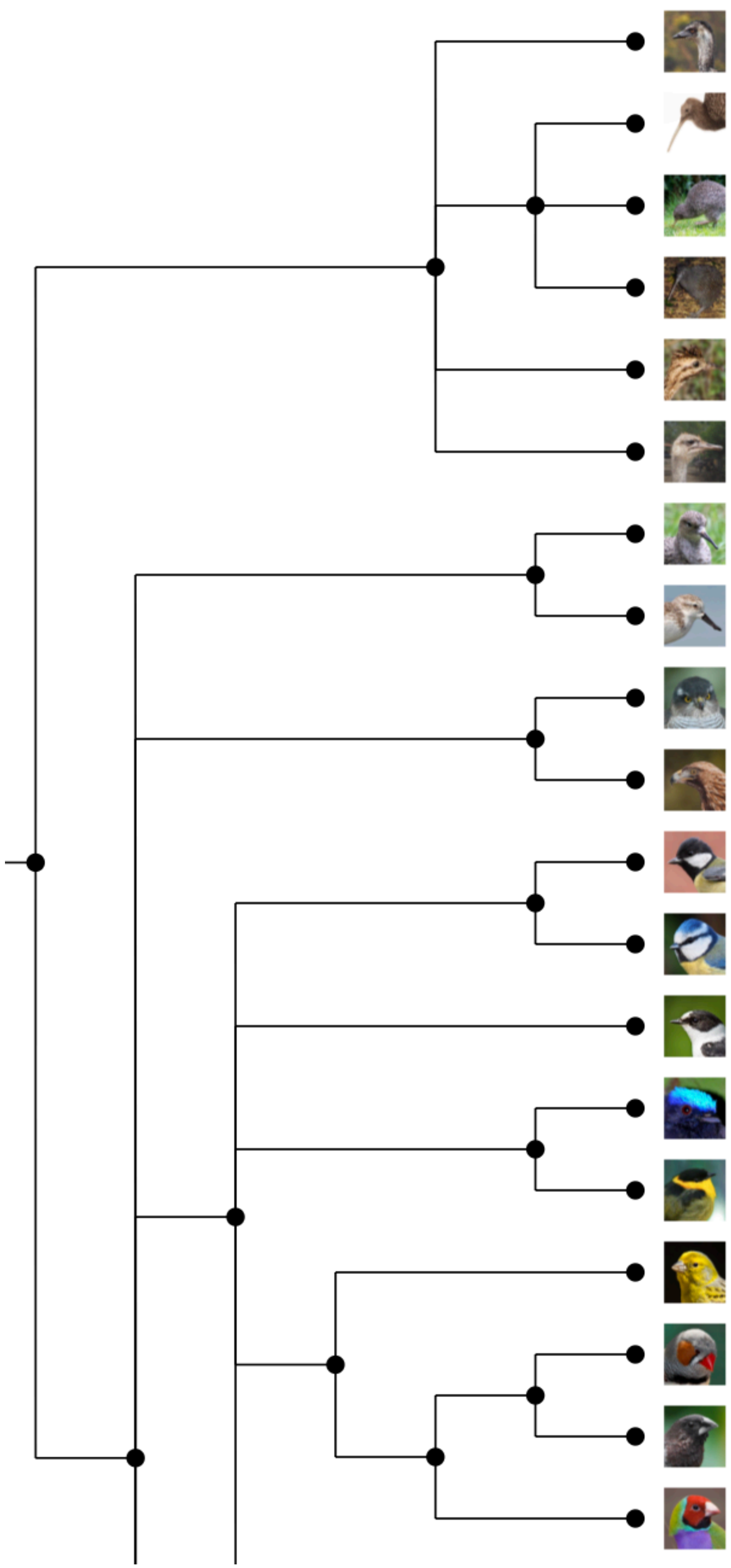

| Animal Name | Protocol (2) - mCSM-PPI2 ΔΔG | Sequence Identity | # Mutated Residue | Sum Grantham Score |
|---|---|---|---|---|
| Emu | 6.607 | 65.83 | 20 | 1449 |
| Great spotted kiwi | 4.371 | 66.17 | 21 | 1613 |
| Little spotted kiwi | 4.521 | 66.29 | 21 | 1613 |
| Okarito brown kiwi | 5.151 | 66.42 | 21 | 1613 |
| Chilean tinamou | 2.884 | 63.81 | 19 | 1256 |
| African ostrich | 4.439 | 62.27 | 20 | 1484 |
| Ruff | 5.231 | 63.67 | 22 | 1573 |
| Spoon-billed sandpiper | 5.232 | 66.00 | 21 | 1431 |
| Eurasian sparrowhawk | 4.157 | 66.42 | 20 | 1299 |
| Golden eagle | 3.902 | 66.00 | 21 | 1240 |
| Great Tit | 2.688 | 67.37 | 22 | 1330 |
| Blue tit | 1.267 | 67.37 | 20 | 1193 |
| Flycatcher | 6.546 | 66.58 | 20 | 1186 |
| Blue-crowned manakin | 4.996 | 66.25 | 23 | 1507 |
| Golden-collared manakin | 5.898 | 65.84 | 23 | 1507 |
| Common canary | 3.786 | 64.89 | 19 | 1127 |
| Zebra finch | 7.275 | 65.34 | 20 | 1182 |
| Bengalese finch | 3.109 | 64.32 | 20 | 1174 |
| Gouldian finch | 2.603 | 66.54 | 21 | 1304 |

### Birds (2)

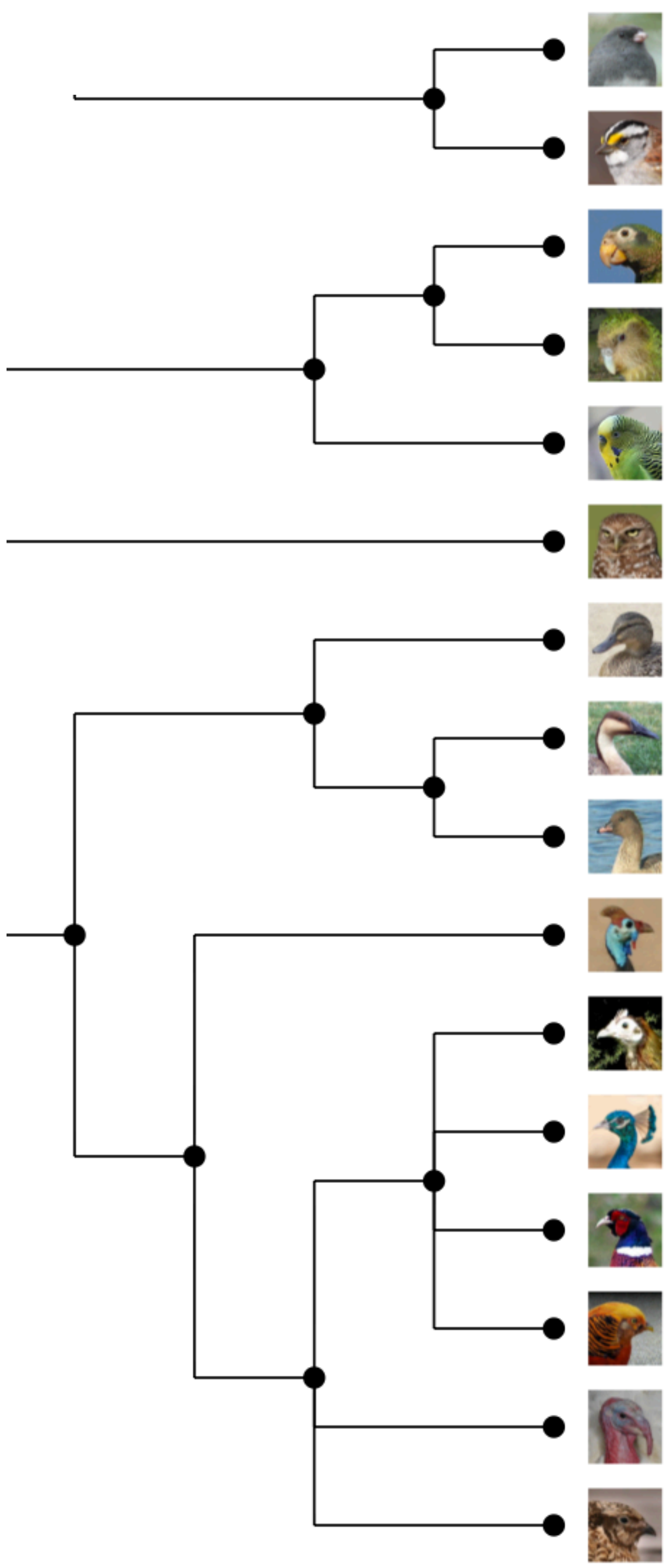

| Animal Name | Protocol (2) - mCSM-PPI2 ΔΔG | Sequence Identity | # Mutated Residue | Sum Grantham Score |
|---|---|---|---|---|
| Dark-eyed junco | 5.861 | 66.46 | 20 | 1338 |
| White-throated sparrow | 4.551 | 65.88 | 21 | 1381 |
| Yellow-billed parrot | 4.227 | 66.50 | 20 | 1376 |
| Kakapo | 6.433 | 61.78 | 21 | 1278 |
| Budgerigar | 3.885 | 62.67 | 20 | 1343 |
| Burrowing owl | 5.064 | 65.79 | 19 | 1287 |
| Duck | 5.889 | 69.29 | 18 | 1394 |
| Swan goose | 3.604 | 65.00 | 17 | 1173 |
| Pink-footed goose | 5.428 | 65.06 | 19 | 1437 |
| Helmeted guineafowl | 5.791 | 65.35 | 20 | 1439 |
| Chicken | 5.001 | 65.97 | 19 | 1350 |
| Indian peafowl | NIL | 55.84 | NIL | NIL |
| Ring-necked pheasant | 5.694 | 65.26 | 19 | 1414 |
| Golden pheasant | NIL | 62.41 | NIL | NIL |
| Turkey | 5.731 | 64.20 | 19 | 1412 |
| Japanese quail | 4.377 | 66.79 | 20 | 1490 |

### Reptiles / Amphibian

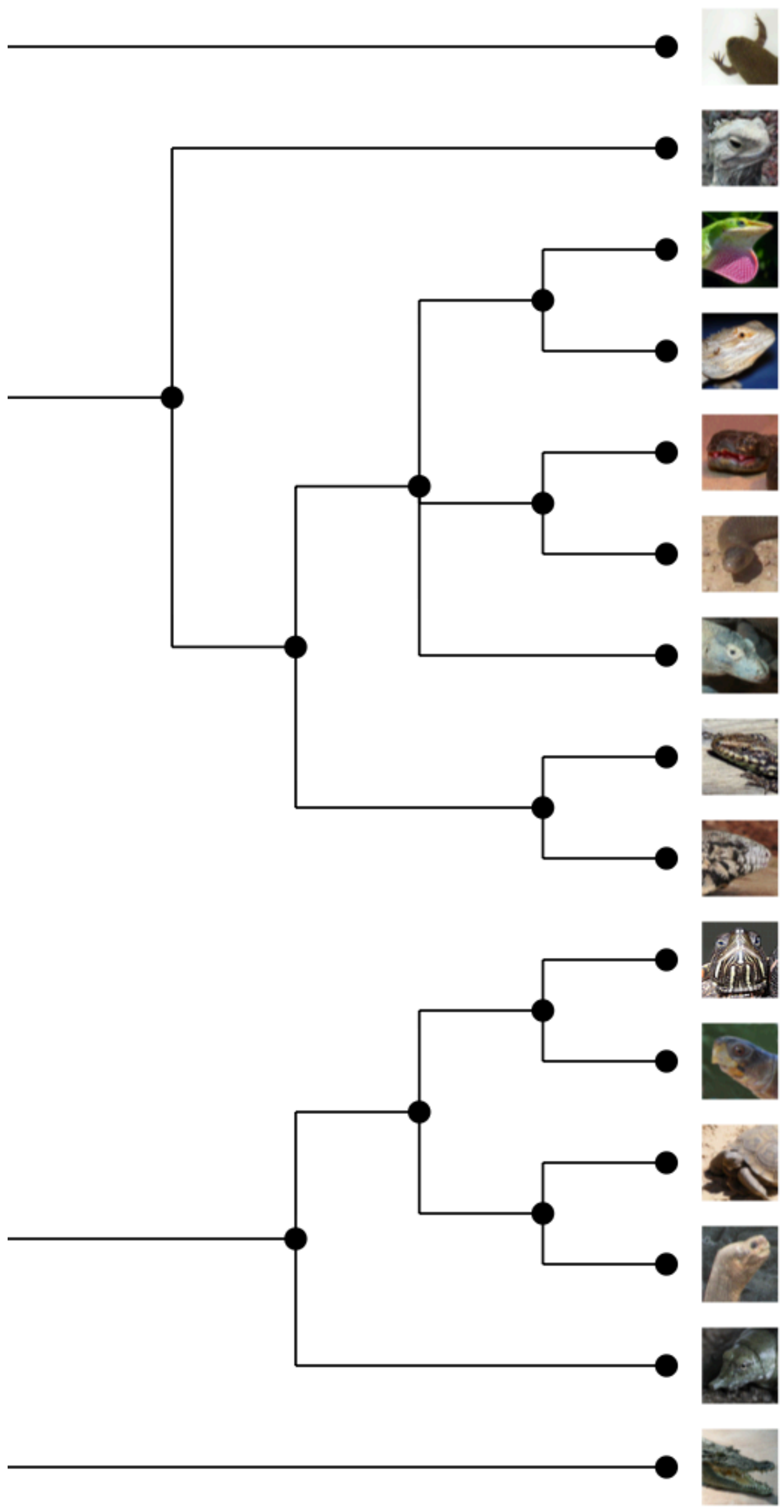

| Animal Name | Protocol (2) - mCSM-PPI2 ΔΔG | Sequence Identity | # Mutated Residue | Sum Grantham Score |
|---|---|---|---|---|
| Tropical clawed frog | 3.883 | 60.15 | 22 | 1458 |
| Tuatara | 2.656 | 64.04 | 16 | 1232 |
| Anole lizard | 5.411 | 63.66 | 22 | 1567 |
| Central bearded dragon | 4.993 | 64.87 | 21 | 1525 |
| Mainland tiger snake | 8.032 | 61.61 | 20 | 1338 |
| Eastern brown snake | 5.643 | 59.50 | 20 | 1280 |
| Komodo dragon | 5.32 | 61.31 | 22 | 1731 |
| Common wall lizard | NIL | 59.09 | NIL | NIL |
| Argentine black and white tegu | 6.117 | 65.05 | 23 | 1501 |
| Painted turtle | 3.616 | 66.46 | 21 | 1583 |
| Three-toed box turtle | 4.223 | 66.96 | 21 | 1583 |
| Agassiz's desert tortoise | NIL | 62.27 | NIL | NIL |
| Abingdon island giant tortoise | 4.756 | 66.75 | 22 | 1605 |
| Chinese softshell turtle | 4.48 | 67.12 | 21 | 1556 |
| Australian saltwater crocodile | 2.533 | 65.63 | 21 | 1463 |

### Fishes (1) and Shark

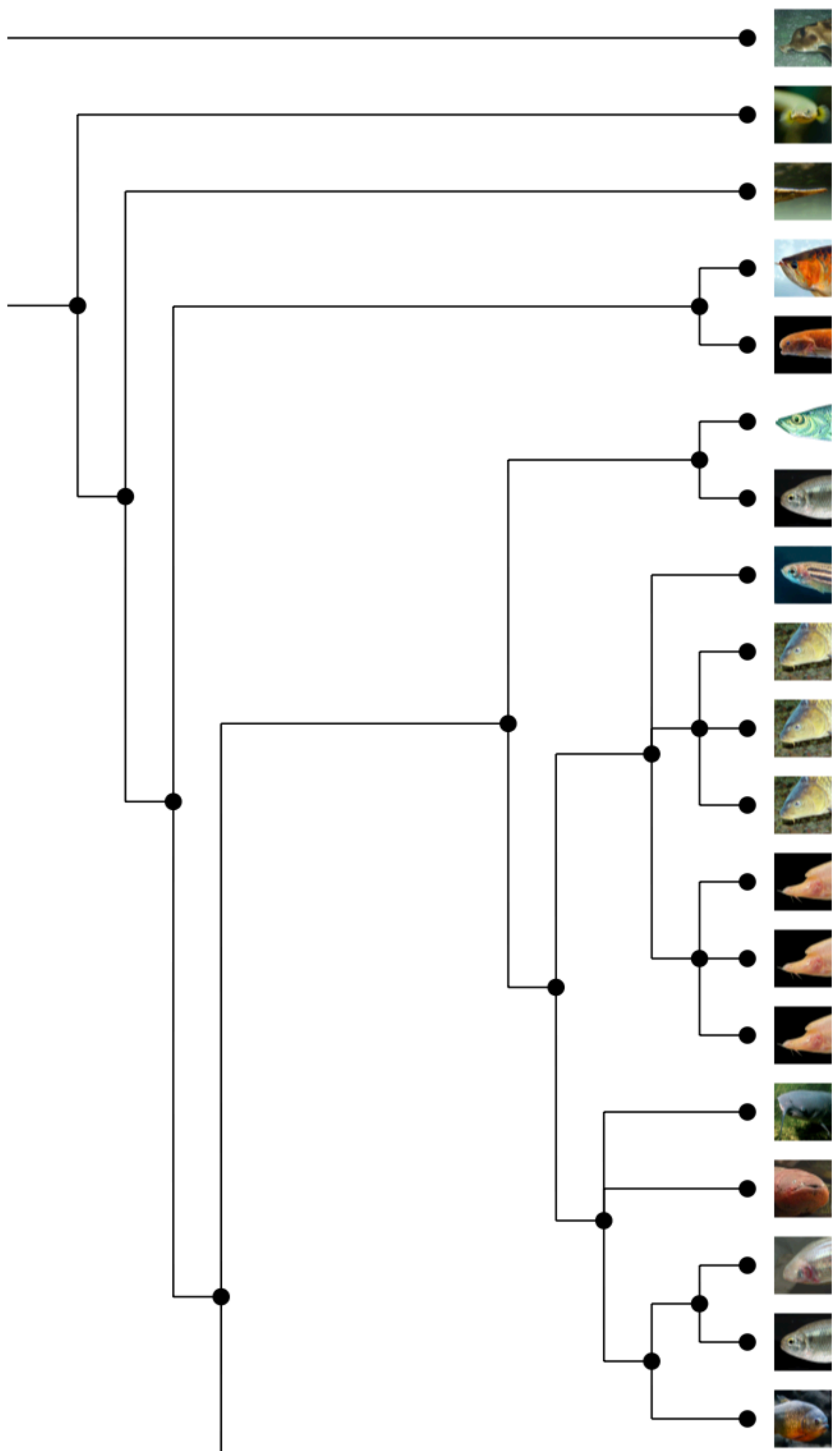

| Animal Name | Protocol (2) - mCSM-PPI2 ΔΔG | Sequence Identity | # Mutated Residue | Sum Grantham Score |
|---|---|---|---|---|
| Elephant shark | 8.569 | 58.96 | 24 | 1524 |
| Reedfish | 6.979 | 57.92 | 18 | 1542 |
| Spotted gar | 5.158 | 58.78 | 25 | 1926 |
| Asian bonytongue | 7.478 | 59.23 | 26 | 1813 |
| Paramormyrops kingsleyae | NIL | NIL | NIL | NIL |
| Atlantic herring | 7.277 | 57.91 | 28 | 1877 |
| Denticle herring | 7.706 | 59.51 | 23 | 1575 |
| Zebrafish | 5.835 | 57.46 | 22 | 1333 |
| Common carp hebaored | 6.242 | 56.34 | 23 | 1501 |
| Common carp huang he | 2.423 | 59.09 | 22 | 1475 |
| Common carp german mirror | 5.798 | 57.57 | 23 | 1563 |
| Golden-line barbel | NIL | NIL | NIL | NIL |
| Horned golden-line barbel | 5.435 | 57.16 | 23 | 1493 |
| Blind barbel | 3.868 | 56.45 | 23 | 1461 |
| Channel catfish | 6.25 | 57.53 | 24 | 1499 |
| Electric eel | 5.731 | 59.07 | 22 | 1499 |
| Pachon cavefish | NIL | NIL | NIL | NIL |
| Mexican tetra | 4.348 | 59.46 | 22 | 1656 |
| Red-bellied piranha | NIL | NIL | NIL | NIL |

### Fishes (2)

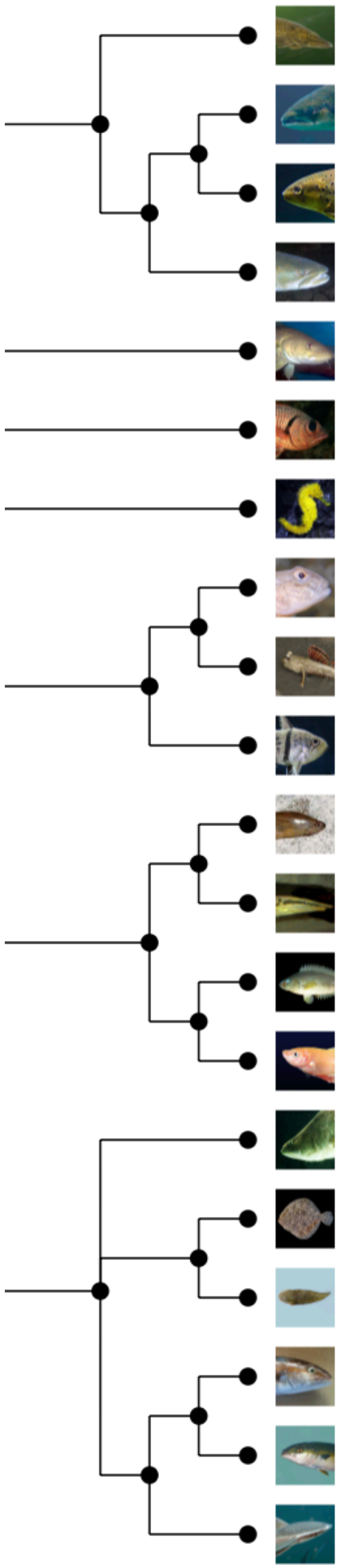

| Animal Name | Protocol (2) - mCSM-PPI2 ΔΔG | Sequence Identity | # Mutated Residue | Sum Grantham Score |
|---|---|---|---|---|
| Northern pike | 2.123 | 58.04 | 26 | 1865 |
| Atlantic salmon | NIL | NIL | NIL | NIL |
| River trout | 4.757 | 57.86 | 26 | 1733 |
| Huchen | 6.074 | 56.22 | 23 | 1507 |
| Cod | 3.852 | 60.03 | 24 | 1597 |
| Pinecone soldierfish | 9.666 | 58.82 | 21 | 1566 |
| Tiger tail seahorse | NIL | NIL | NIL | NIL |
| Round goby | 9.83 | 56.65 | 27 | 1644 |
| Periophthalmus magnuspinnatus | NIL | NIL | NIL | NIL |
| Orbiculate cardinalfish | NIL | NIL | NIL | NIL |
| Swamp eel | 4.295 | 54.24 | 22 | 1596 |
| Zig-zag eel | 6.014 | 57.91 | 26 | 1753 |
| Climbing perch | 8.16 | 58.96 | 26 | 1648 |
| Siamese fighting fish | 3.956 | 58.89 | 22 | 1521 |
| Barramundi perch | 3.028 | 58.86 | 22 | 1453 |
| Turbot | 3.218 | 57.36 | 23 | 1607 |
| Tongue sole | 9.281 | 57.56 | 23 | 1489 |
| Greater amberjack | NIL | NIL | NIL | NIL |
| Yellowtail amberjack | 7.959 | 58.15 | 24 | 1619 |
| Live sharksucker | 7.637 | 57.97 | 25 | 1738 |

### Fishes (3)

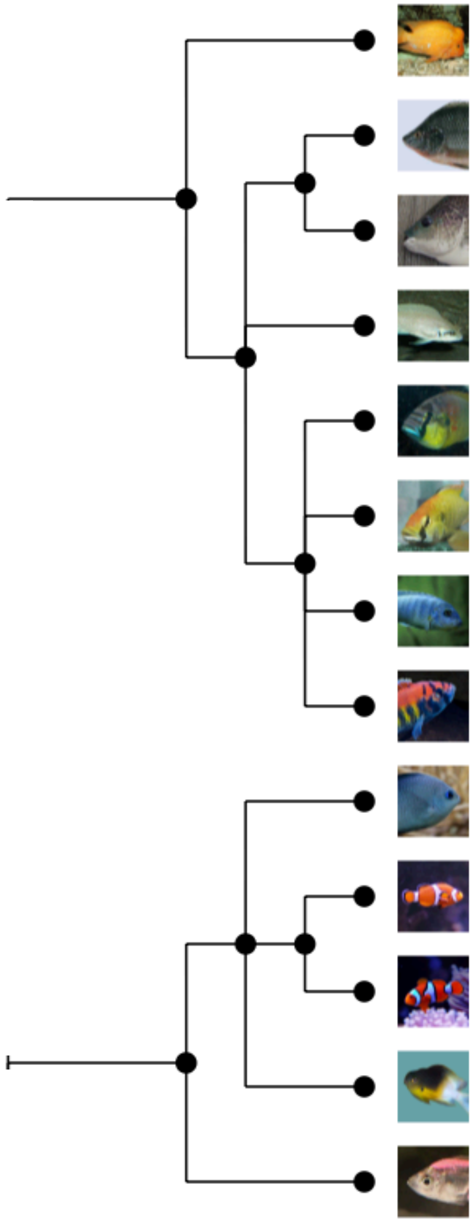

| Animal Name | Protocol (2) - mCSM-PPI2 $\Delta\Delta G$ | Sequence Identity | # Mutated Residue | Sum Grantham Score |
|---|---|---|---|---|
| Midas cichlid | 2.188 | 57.69 | 23 | 1538 |
| Nile tilapia | 0.664 | 56.74 | 20 | 1364 |
| Blue tilapia | 3.864 | 57.16 | 21 | 1387 |
| Lyretail cichlid | 3.149 | 58.07 | 21 | 1432 |
| Burton's mouthbrooder | 3.556 | 58.07 | 21 | 1387 |
| Eastern happy | 3.47 | 58.07 | 21 | 1387 |
| Zebra mbuna | 4.075 | 58.07 | 21 | 1387 |
| Makobe Island cichlid | 4.95 | 58.89 | 21 | 1387 |
| Spiny chromis | 5.419 | 56.47 | 26 | 1736 |
| Clown anemonefish | 7.337 | 58.18 | 23 | 1755 |
| Orange clownfish | 5.599 | 56.34 | 23 | 1755 |
| Bicolor dameselfish | NIL | 64.32 | NIL | NIL |
| Indian glassy fish | 6.815 | 56.27 | 24 | 1744 |

### Fishes (4)

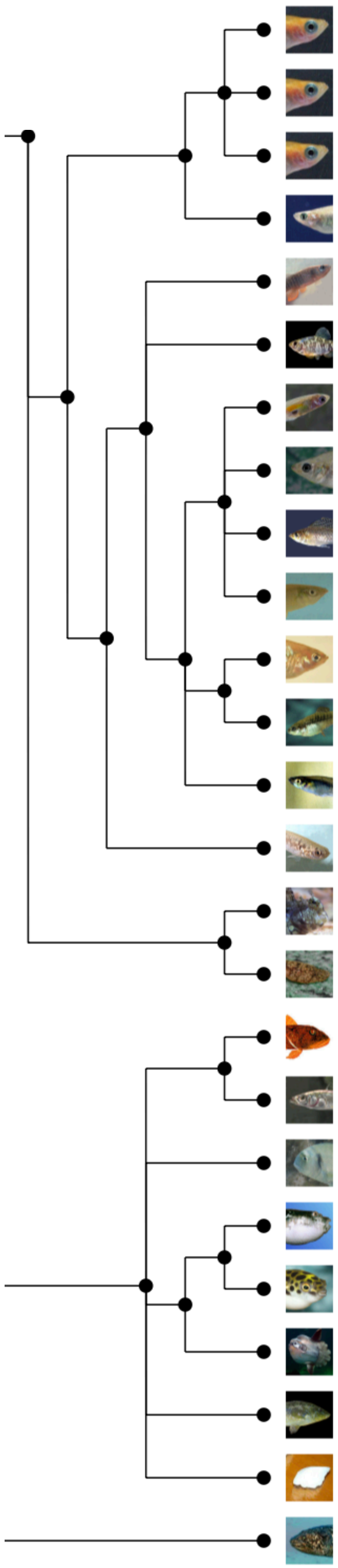

| Animal Name | Protocol (2) - mCSM-PPI2 ΔΔG | Sequence Identity | # Mutated Residue | Sum Grantham Score |
|---|---|---|---|---|
| Japanese medaka HSOK | 7.204 | 57.98 | 24 | 1817 |
| Japanese medaka HdrR | NIL | NIL | NIL | NIL |
| Japanese medaka HNI | 5.987 | 56.94 | 24 | 1817 |
| Indian medaka | 3.99 | 55.84 | 24 | 1802 |
| Mummichog | 7.821 | 58.36 | 24 | 1531 |
| Sheepshead minnow | 4.826 | 58.99 | 24 | 1565 |
| Guppy | NIL | NIL | NIL | NIL |
| Amazon molly | 5.66 | 57.53 | 25 | 1617 |
| Sailfin molly | NIL | NIL | NIL | NIL |
| Shortfin molly | 6.512 | 57.65 | 25 | 1617 |
| Platyfish | 6.332 | 58.60 | 25 | 1649 |
| Monterrey platyfish | NIL | NIL | NIL | NIL |
| Western mosquitofish | 8.745 | 58.03 | 25 | 1649 |
| Mangrove rivulus | 6.228 | 58.33 | 23 | 1763 |
| Jewelled blenny | 6.587 | 56.97 | 22 | 1640 |
| Blunt-snouted clingfish | 8.498 | 59.87 | 26 | 1835 |
| Channel bull blenny | 6.876 | 56.80 | 22 | 1659 |
| Stickleback | 4.767 | 58.11 | 22 | 1659 |
| Gilthead seabream | 8.471 | 58.23 | 23 | 1480 |
| Fugu | 5.871 | 55.31 | 23 | 1684 |
| Tetraodon | 5.336 | 57.20 | 24 | 1814 |
| Ocean sunfish | 5.064 | 58.85 | 23 | 1633 |
| Ballan wrasse | 8.614 | 56.18 | 25 | 1891 |
| Large yellow croaker | 3.538 | 58.99 | 23 | 1435 |
| Coelacanth | 7.845 | 60.96 | 25 | 1744 |
