## Supplementary Materials for "SARS-CoV-2 spike protein predicted to form complexes with host receptor protein orthologues from a broad range of mammals"

#### Supplementary Results

##### Supplementary Results 1: Conservation of ACE2 in vertebrates

The structural models were compared to the human ACE2 protein structure using SSAP[1]. Most models had high structural similarity to human ACE2 (SSAP > 90) (Supplementary Fig. 1B and Supplementary Fig. 1C).

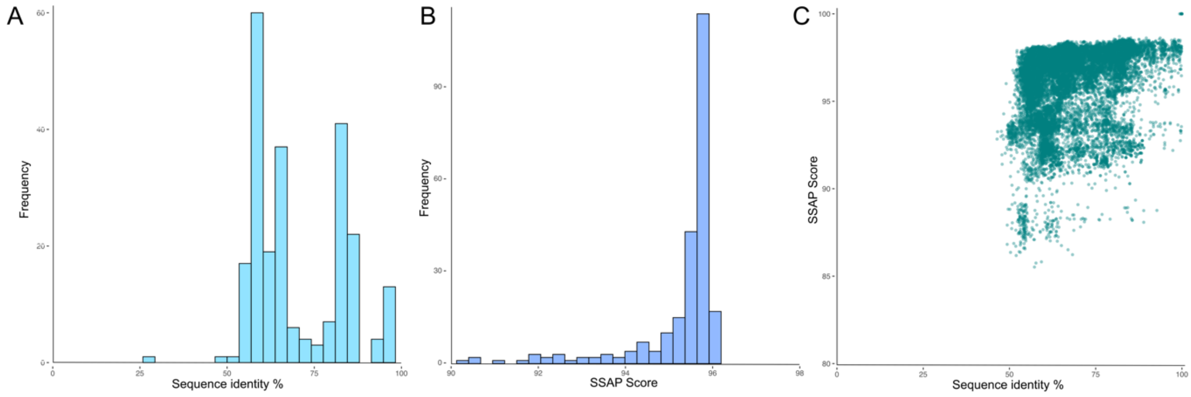


***Supplementary Figure 1: Conservation of ACE2. (a) ACE2 sequence conservation in vertebrates. Distribution of pairwise sequence identities of ACE2 orthologues compared to human ACE2. (b) ACE2 structure conservation in vertebrates. Distribution of pairwise SSAP scores of ACE2 orthologues compared to human ACE2. (c) Pairwise sequence identities and SSAP scores for ACE2 orthologues.***

However, despite sharing > 60% sequence identity to the human ACE2, some models had lower structural similarity (Fig. 1C). These changes probably reflect mutations slightly altering the relative orientations of the S-protein and ACE2 across the interface. (Supplementary Fig. 2).

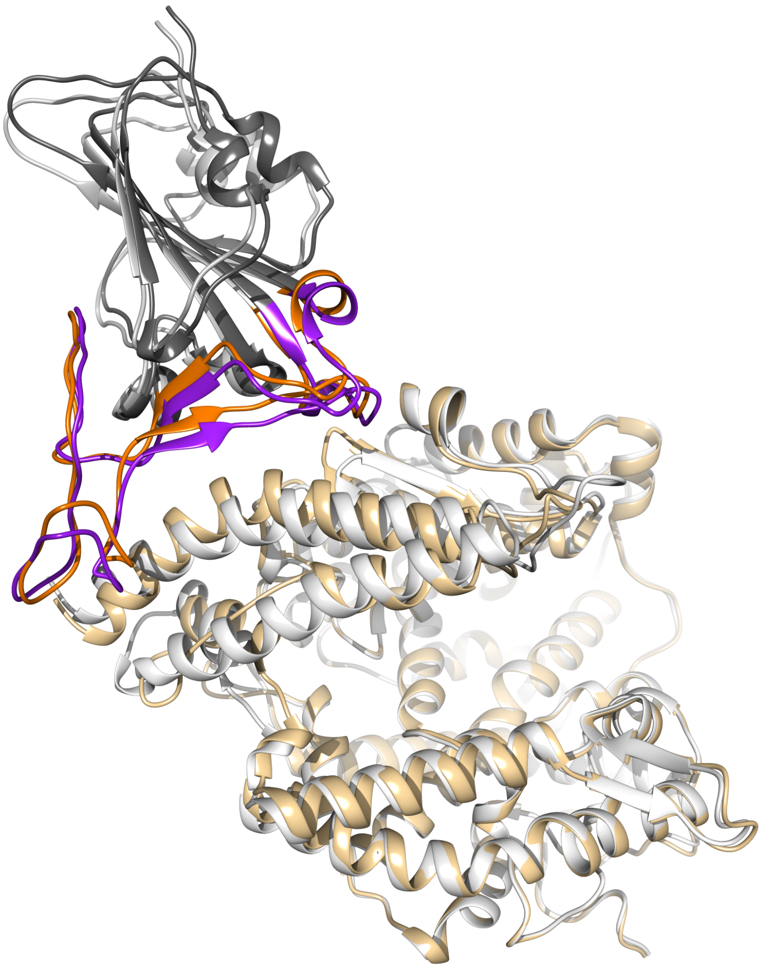

***Supplementary Figure 2: Superposition of the S-protein:ACE2 complexes from human and the model from climbing perch (Anabas testudineus). There is 59.5% sequence identity in ACE2 sequences of human and climbing perch.***

#### Supplementary Results 2: Identification of critical S-protein:ACE2 interface residues

ACE2 residues directly contacting the S-protein were identified in a structure of the complex. We also identified a more extended set of both DC residues and residues within 8A of DC residues likely to be influencing binding. Residues within 5Å of the interacting protein are reported in PDBe and PDBsum[2,3], and we refer to these as 'Direct Contact' (DC) residues. Other important residues likely to influence binding have been identified using a variety of approaches, including structural analysis by us and other groups, alanine scanning, and mutagenesis[4–10]. In addition, we identified highly conserved residues, sites under positive selection and predicted allosteric sites. This larger set includes 42 residues that are not involved in direct contact with the interacting protein but could still influence the binding affinity. 15 of these residues are within 5Å and 27 residues are further away but within 8Å of one, or more, DC residues. We, therefore, compiled a set of residues that includes these residues, together with the DC residues. We refer to this as the direct contact plus 'extended set' of residues ('DCEX' residues) (Fig. 1b). Some of the ACE2 DCEX residues are highly-conserved positions in the ACE2 FunFam (see Supplementary Fig. 3).

| Residue | Residue type | Structure analyses | Deep mutagenesis | Computational Alanine Scanning | Evidence of positive selection | Potential allosteric region in our study | Allosteric site prediction by PARS/ENM/ AllositePro | Computational ACE2-based peptide inhibitor design | ScoreCons |
|---|---|---|---|---|---|---|---|---|---|
| GLN24 | DC | ■ | | ■ | ■ | | | ■ | 0.245 |
| THR27 | DC | ■ | | | ■ | | | ■ | 0.289 |
| PHE28 | DC | ■ | | | ■ | | | ■ | 0.583 |
| ASP30 | DC | ■ | | | ■ | | | ■ | 0.307 |
| LYS31 | DC | ■ | | | ■ | | | ■ | 0.338 |
| HIS34 | DC | ■ | | | ■ | | | ■ | 0.234 |
| GLU35 | DC | ■ | | | ■ | | | ■ | 0.397 |
| GLU37 | DC | ■ | | ■ | ■ | | | ■ | 0.353 |
| ASP38 | DC | ■ | | ■ | ■ | | | ■ | 0.358 |
| TYR41 | DC | ■ | | ■ | | | | ■ | 0.538 |
| GLN42 | DC | ■ | | ■ | ■ | | | ■ | 0.440 |
| LEU79 | DC | ■ | | | ■ | | | ■ | 0.253 |
| MET82 | DC | ■ | | | | | | ■ | 0.295 |
| TYR83 | DC | ■ | | ■ | | | | ■ | 0.546 |
| ASN330 | DC | ■ | | | | | | ■ | 0.741 |
| LYS353 | DC | ■ | | ■ | | | | ■ | 0.362 |
| GLY354 | DC | ■ | | | ■ | | ■ | ■ | 0.475 |
| ASP355 | DC | ■ | | ■ | | | ■ | ■ | 0.918 |
| ARG357 | DC | ■ | | ■ | | | | ■ | 0.894 |
| ARG393 | DC | ■ | | | ■ | | | ■ | 0.89 |
| SER19 | 5A | ■ | | | | | | ■ | 0.013 |
| LEU45 | 5A | ■ | | | ■ | | | ■ | 0.563 |
| GLU329 | 5A | ■ | | | ■ | ■ | ■ | ■ | 0.469 |
| LEU29 | 5A | | ■ | | ■ | | | ■ | 0.583 |
| PHE72 | 5A | | ■ | | | | | ■ | 0.542 |
| THR324 | 5A | | ■ | | | ■ | ■ | ■ | 0.556 |
| TRP69 | 8A | | ■ | | ■ | | | ■ | 0.536 |
| GLN325 | 8A | | ■ | | | ■ | ■ | ■ | 0.521 |
| LYS26 | 5A | | ■ | | ■ | | ■ | ■ | 0.337 |
| LEU39 | 5A | | ■ | | | | | ■ | 0.417 |
| PHE40 | 5A | | ■ | | | | | ■ | 0.280 |
| GLU75 | 5A | | ■ | | | | | ■ | 0.396 |
| GLN76 | 5A | | ■ | | | | | ■ | 0.309 |
| ALA386 | 5A | | ■ | | | | ■ | ■ | 0.546 |
| PRO389 | 5A | | ■ | | | | | ■ | 0.670 |
| GLY352 | 5A | | | ■ | | ■ | ■ | ■ | 0.819 |
| PHE356 | 5A | | | ■ | | ■ | ■ | ■ | 0.815 |
| PHE390 | 5A | | | ■ | | | | ■ | 0.715 |
| TYR381 | 8A | | | ■ | | | | ■ | 0.900 |
| TRP328 | 5A | | | | | ■ | ■ | ■ | 0.910 |
| SER331 | 5A | | | | | ■ | | ■ | 0.929 |
| MET332 | 5A | | | | | ■ | | ■ | 0.914 |
| THR347 | 5A | | | | | ■ | | ■ | 0.923 |
| ALA348 | 5A | | | | | ■ | ■ | ■ | 0.927 |
| TRP349 | 5A | | | | | ■ | | ■ | 0.919 |
| ASP350 | 5A | | | | | ■ | | ■ | 0.919 |
| ILE358 | 5A | | | | | ■ | | ■ | 0.908 |
| LEU359 | 5A | | | | ■ | ■ | | ■ | 0.711 |
| TYR385 | 5A | | | | | | | ■ | 0.905 |
| LEU391 | 5A | | | | | | | ■ | 0.774 |
| GLY395 | 5A | | | | | | ■ | ■ | 0.907 |
| LEU333 | 8A | | | | | ■ | | ■ | 0.779 |
| LEU560 | 8A | | | | | | | | 0.804 |
| ALA384 | 8A | | | | | | ■ | ■ | 0.814 |
| SER563 | 8A | | | | | | | | 0.849 |
| MET323 | 8A | | | | | ■ | ■ | ■ | 0.867 |
| GLU375 | 8A | ■ | | | | | | ■ | 0.903 |
| ALA396 | 8A | | | | | | | ■ | 0.905 |
| MET383 | 8A | | | | | | ■ | ■ | 0.906 |
| MET360 | 8A | | | | | ■ | | ■ | 0.907 |
| HIS401 | 8A | | | | | | | | 0.908 |

***Supplementary Figure 3: Table of direct contact (DC) residues in ACE2 that are involved in direct contact with the S-protein, and direct contact extended (DCEX) residues in ACE2 that are within 8Å of DC residues. We indicate whether residues are in direct contact, within 5Å or between 5Å and 8Å from the S-protein. We list all structural evidence that we have for each residue, including structure analyses, deep mutagenesis, computational alanine scanning, evidence of positive selection, potential allosteric region, allosteric site prediction, computational ACE-based peptide inhibitor design (Han Y, Král P 2020) and ScoreCons scores. The higher the ScoreCons value the more conserved the residue, as indicated by a deeper red colour.***

#### Supplementary Results 3: Changes in the energy of the S-protein:ACE2 complex in vertebrates

We used two protocols, protocols 1 and 2, to assess the relative change in binding energy (ΔΔG) of the SARS-CoV-2 S-protein:ACE2 complex following mutations in the DC and DCEX residues. Since these residues lie directly in the binding interface (DC residues) or in the secondary shell and are likely to be influencing binding, we hypothesise that the changes in energy of the complex caused by mutations in these residues can be used to gauge the relative risk of infection for each animal. We examined how well these approaches correlated with each other (Supplementary Fig. 4). We also correlated the ΔΔG values (protocols 1 and 2) with SARS-CoV-2 infection phenotypes in different animals[4,11–17] (Table 1; phenotype data are sourced from in vivo[12,13,16,18,19] and in vitro[4,14,15] infection studies, some of which have not been peer reviewed).

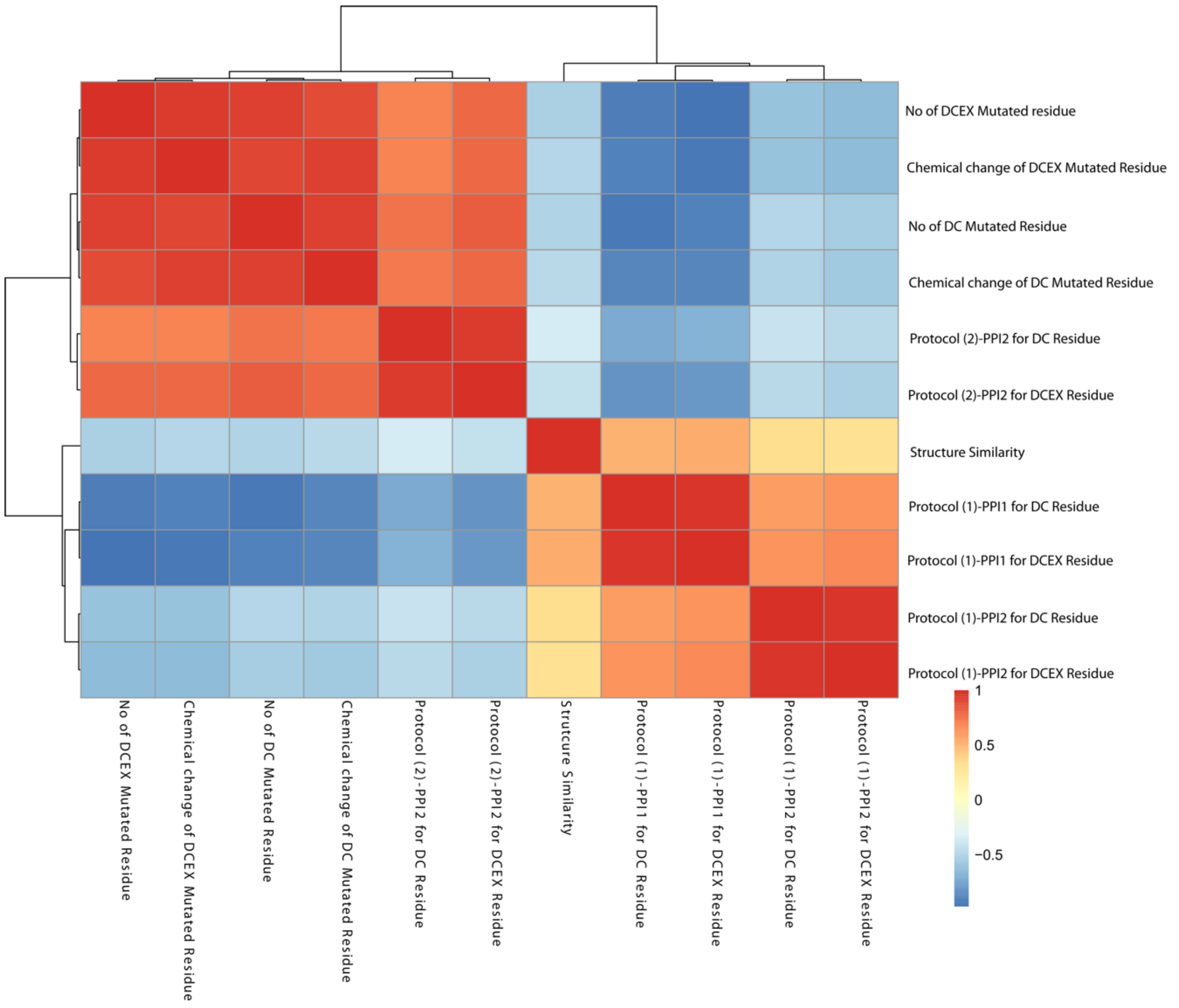


***Supplementary Figure 4: Heatmap showing the Spearman rank correlation coefficients between the number of mutated residues (DC and DCEX), chemical change of the mutated residues, as calculated by the Grantham score (DC and DCEX), Protocol (1)-PPI1 and PPI2 scores (DC and DCEX), Protocol (2)-PPI2 scores (DC and DCEX), and SSAP structure similarity scores.***

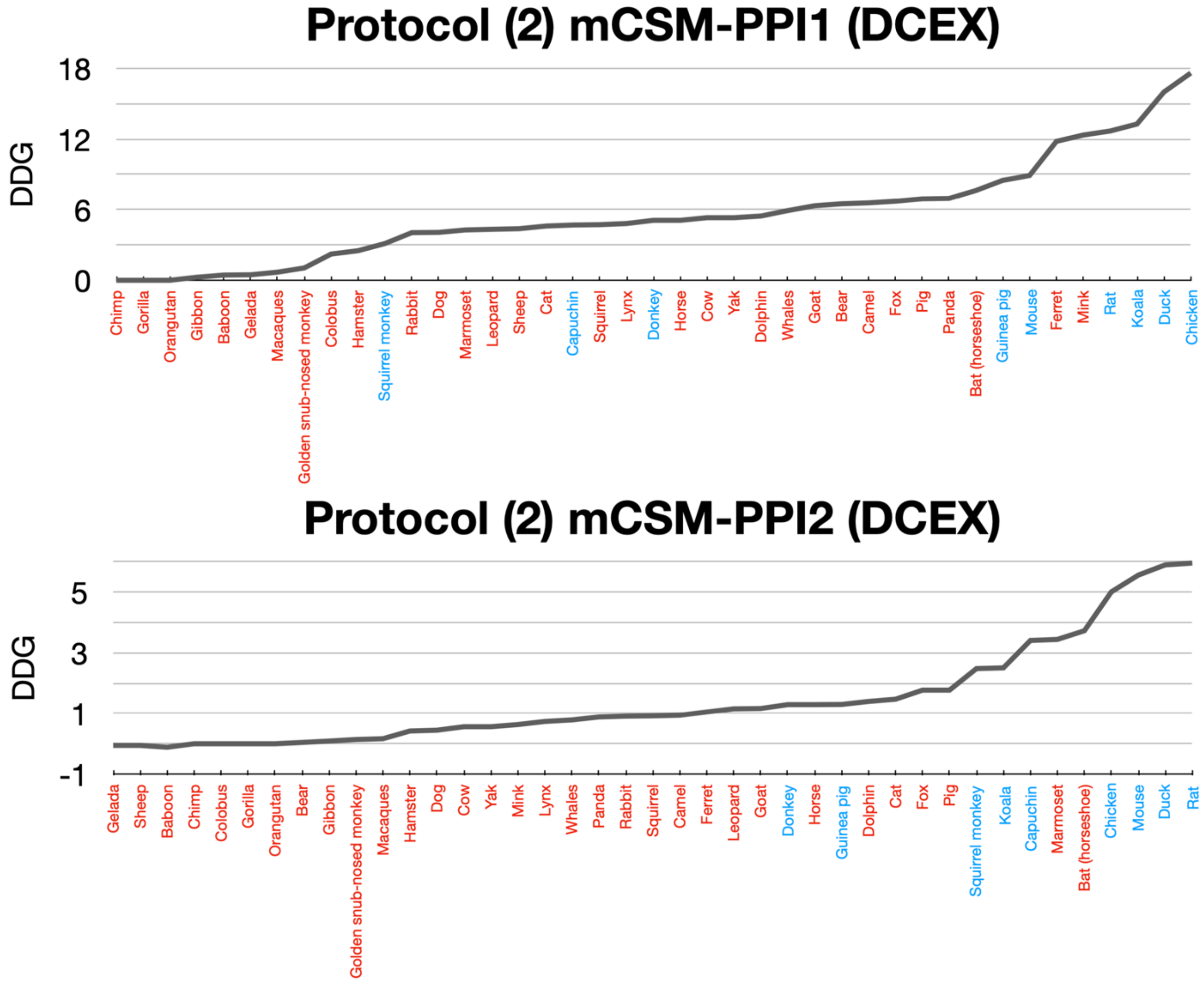


***Supplementary Figure 5: Anti-correlation plot of mCSM-PPI2 values from protocol 1 and 2. Experimental data suggests that the animals in red are at risk of infection whilst the animals in blue are not.***

In order to further verify our predictions, we also applied another independent method HADDOCK[20] (see Supplementary Methods 4) to the 41 animals for which we have experimental evidence. In Supplementary Figure 6 we plot the correspondence between mCSM-PPI2 and HADDOCK values. By setting the threshold for animals at risk using the experimental data (i.e. a threshold of ΔΔG 3.7 for mCSM-PPI2 and a threshold of -136 a.u for HADDOCK, animals at risk are those points within the box formed by the dotted lines on the plot) we can see considerable agreement between the two methods with agreement for nearly 95% of the animals predicted to be at risk.

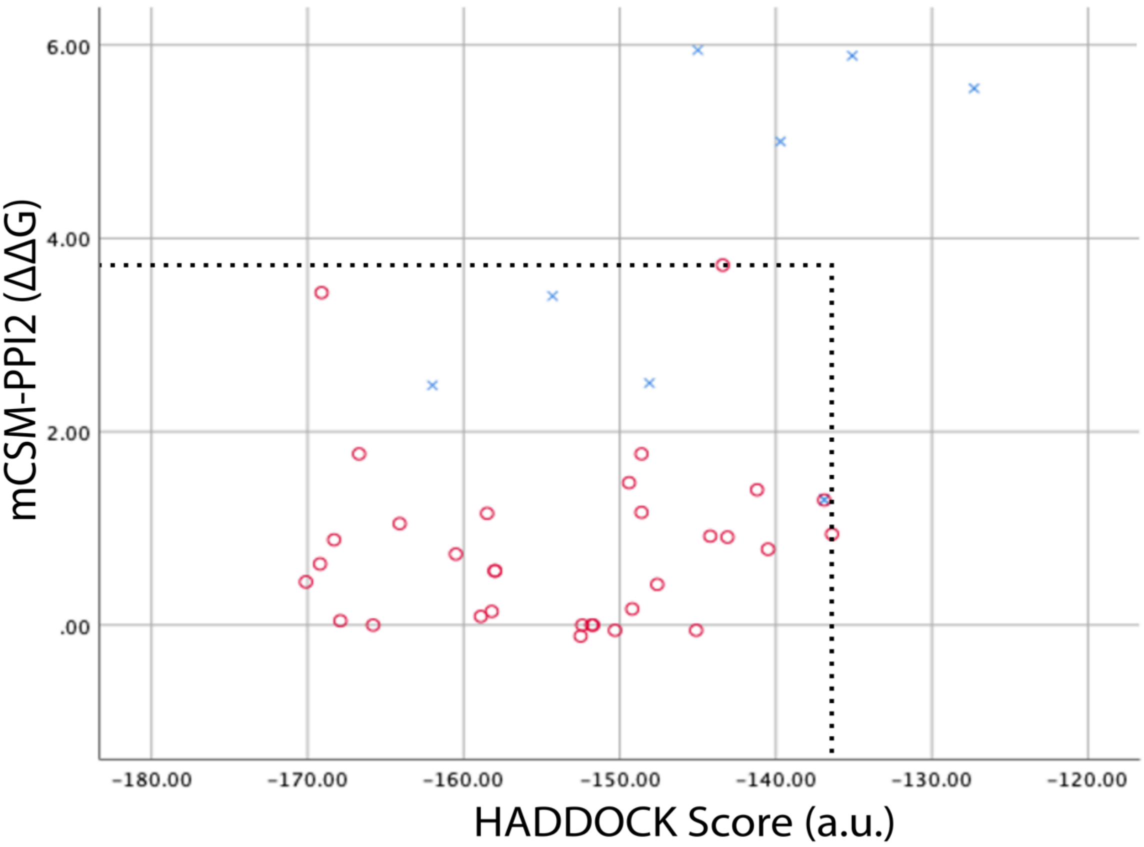


***Supplementary Figure 6: Plot of mCSM-PPI2 ΔΔG versus HADDOCK a.u. score for 41 animals with experimental evidence. Red denotes animals for which experimental studies found evidence of infection. Blue denotes animals for which experimental studies found no evidence of infection.***

For Supplementary Fig. 7, please refer to the accompanying PDF.

***Supplementary Figure 7: Multiple phylogenetic trees, for different taxonomic levels, for all vertebrates species that were analysed, showing changes in the energy of the S-protein:ACE2 complex, number of mutated residues and sum Grantham scores. Animals are categorised according to risk of infection by SARS-CoV-2, with ΔΔG ≤ 3.7 high risk (red), and ΔΔG > 3.7 low risk (blue). These thresholds were chosen as they agree well with the available experimental data. Animal photos courtesy of ENSEMBL and associated sources (https://www.ensembl.org/info/about/image_credits.html)***

Below we show the residues that P(2)-PPI2 reports as stabilising or destabilising for the SARS-CoV-2 S-protein:ACE2 animal complex for DC (Supplementary Fig. 8) and DCEX (Supplementary Fig. 9) residues.

| | 24 | 27 | 28 | 30 | 31 | 34 | 35 | 37 | 38 | 41 | 42 | 79 | 82 | 83 | 330 | 353 | 354 | 355 | 357 | 393 |
|---|---|---|---|---|---|---|---|---|---|---|---|---|---|---|---|---|---|---|---|---|
| Human | Q | T | F | D | K | H | E | E | D | Y | Q | L | M | Y | N | K | G | D | R | R |
| Bonobo | Q | T | F | D | K | H | E | E | D | Y | Q | L | M | Y | N | K | G | D | R | R |
| Macaque | Q | T | F | D | K | H | E | E | D | Y | Q | L | M | Y | N | K | G | D | R | R |
| Orangutan | Q | T | F | D | K | H | E | E | D | Y | Q | L | M | Y | N | K | G | D | R | R |
| Chimpanzee | Q | T | F | D | K | H | E | E | D | Y | Q | L | M | Y | N | K | G | D | R | R |
| Golden Snub-nosed Monkey | Q | T | F | D | K | H | E | E | D | Y | Q | L | M | Y | N | K | G | D | R | R |
| Gorilla | Q | T | F | D | K | H | E | E | D | Y | Q | L | M | Y | N | K | G | D | R | R |
| Bushbaby | Q | T | F | D | N | R | E | E | E | H | Q | I | T | Y | N | K | D | D | R | R |
| Mouse | N | T | F | N | N | Q | E | E | D | Y | Q | T | S | F | N | H | G | D | R | R |
| Rat | K | S | F | N | K | Q | E | E | D | Y | Q | I | N | F | N | H | G | D | R | R |
| Guinea Pig | Q | T | F | D | E | L | K | E | D | Y | Q | L | A | Y | N | K | N | D | R | R |
| Squirrel | - | T | F | D | K | Q | E | E | D | Y | Q | L | A | Y | N | K | G | D | R | R |
| Chinese Hamster CH0K1GS | Q | T | F | D | K | Q | E | E | D | Y | Q | L | N | Y | N | K | G | D | R | R |
| Golden Hamster | Q | T | F | D | K | Q | E | E | D | Y | Q | L | N | Y | N | K | G | D | R | R |
| Rabbit | - | T | F | E | K | Q | E | E | D | Y | Q | L | T | Y | N | K | G | D | R | R |
| Cow | Q | T | F | E | K | H | E | E | D | Y | Q | M | T | Y | N | K | G | D | R | R |
| Cow Hybrid - Bos Indicus | Q | T | F | E | K | H | E | E | D | Y | Q | M | T | Y | N | K | G | D | R | R |
| Domestic Yak | Q | T | F | E | K | H | E | E | D | Y | Q | M | T | Y | N | K | G | D | R | R |
| Wild Yak | Q | T | F | E | K | H | E | E | D | Y | Q | M | T | Y | N | K | G | D | R | R |
| Sheep | Q | T | F | E | K | H | E | E | D | Y | Q | M | T | Y | N | K | G | D | R | R |
| Goat | Q | T | F | E | K | H | E | E | D | Y | Q | M | T | Y | N | K | G | D | R | R |
| Pig | - | T | F | E | K | L | E | E | D | Y | Q | I | T | Y | N | K | G | D | R | R |
| Arabian Camel | - | T | F | E | E | H | E | E | D | Y | Q | T | T | Y | N | K | G | D | R | R |
| American Mink | - | T | F | E | K | Y | E | E | E | Y | Q | H | T | Y | N | K | H | D | R | R |
| Ferret | - | T | F | E | K | Y | E | E | E | Y | Q | H | T | Y | N | K | R | D | R | R |
| Panda | - | T | F | E | K | Y | E | E | D | Y | Q | H | T | Y | N | K | G | D | R | R |
| Polar Bear | - | T | F | E | K | Y | E | E | D | Y | Q | H | T | Y | N | K | G | D | R | R |
| Red Fox | - | T | F | E | K | Y | E | E | E | Y | Q | L | T | Y | N | K | G | D | R | R |
| Dog | - | T | F | E | K | Y | E | E | E | Y | Q | L | T | Y | N | K | G | D | R | R |
| Cat | - | T | F | E | K | H | E | E | E | Y | Q | L | T | Y | N | K | G | D | R | R |
| Leopard | - | T | F | E | K | H | E | E | E | Y | Q | L | T | Y | N | K | G | D | R | R |
| Donkey | - | T | F | E | K | S | E | E | E | H | Q | L | T | Y | N | K | G | D | R | R |
| Horse | - | T | F | E | K | S | E | E | E | H | Q | L | T | Y | N | K | G | D | R | R |
| Greater Horseshoe Bat | - | K | F | D | D | S | E | E | N | H | Q | L | N | F | N | K | G | D | R | R |
| Hedgehog | E | K | F | D | D | R | Q | E | N | Y | E | T | N | Y | N | N | G | D | R | R |
| Malayan Pangolin | E | T | F | E | K | S | E | E | E | Y | Q | I | N | Y | N | K | H | D | R | R |
| Eurasian Common Shrew | N | K | F | E | N | K | D | E | D | Y | N | I | T | F | N | K | N | D | R | R |
| Masked Palm Civet | - | T | F | E | T | Y | E | Q | E | Y | Q | L | T | Y | N | K | G | D | R | R |
| Elephant | - | T | F | D | T | Q | E | E | D | Y | Q | L | D | F | N | K | G | D | R | R |
| Koala | R | E | F | E | T | K | E | E | E | Y | Q | i | T | F | N | K | G | D | R | R |

X (red) A particular mutation is stabilising the S protein: ACE2 complex
X (blue) The mutation is destabilising the complex
X (grey) Residues identical to human

***Supplementary Figure 8: Direct contact (DC) residues for animals that come into contact with humans. Using mCSM-PPI scores, we show the residues for each animal in contact with humans and indicate whether a particular mutated residue destabilises the S-protein:ACE2 complex in the animal (blue), whether the mutation stabilises the complex (red), and identical in animals and human (grey).***

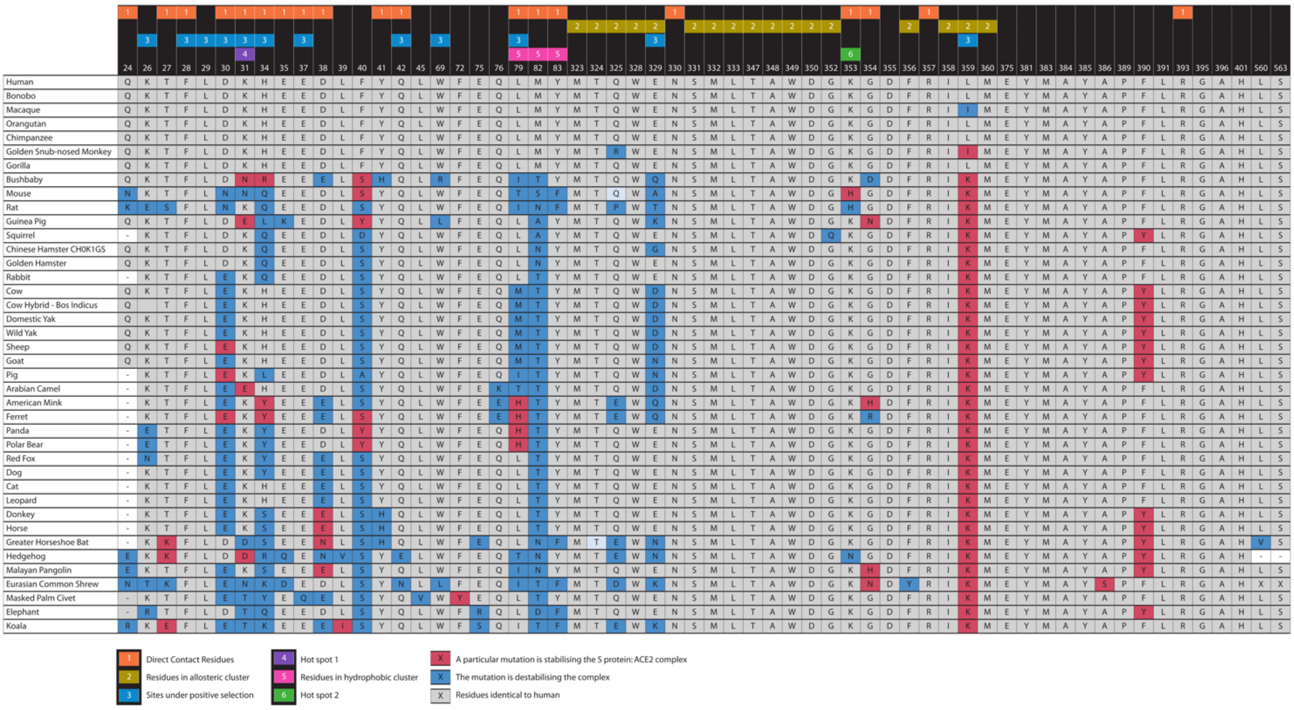


***Supplementary Figure 9: Direct contact extended (DCEX) residues for animals that come into contact with humans. Using mCSM-PPI scores, we show the residues for each animal in contact with humans and indicate whether a particular mutated residue destabilises the S-protein:ACE2 complex in the animal (blue), whether the mutation stabilises the complex (red), and identical in animals and human (grey). Reasons why each residue was selected to be analysed are shown at the top of the table: direct contact, structural analysis, alanine scanning, conservation, allosteric site and/or under positive selection.***

#### Supplementary Results 4: Structural analyses of S-protein:ACE2 interactions

*Cross-species comparison*

We selected twelve animals for structural analyses because they are likely to come into frequent contact with humans in domestic, zoological or agricultural settings (cat, dog, macaque, cow, sheep and camel) or because our energy calculations suggested risk but currently available COVID-19 infection data was ambiguous or suggested no infection (guinea pig, marmoset, capuchin, squirrel monkey, koala) or where we predicted large energy changes despite experimental evidence of infection (horseshoe bat).

Structural variations in the S-protein:ACE2 interface amongst models of these twelve species are highlighted with reference to the human PDB structure (PDB ID 6M0J) and in the context of three sites on the interface: hydrophobic pocket, hotspot-353 and hotspot-31[21] (Supplementary Fig. 10a). These sites have previously been identified [22] as key to understanding why the SARS-CoV-2 S-protein binds to human ACE2 with high affinity and how the viral S-protein has evolved to bind with much higher affinity to human ACE2 than SARS-CoV[5,21,23].

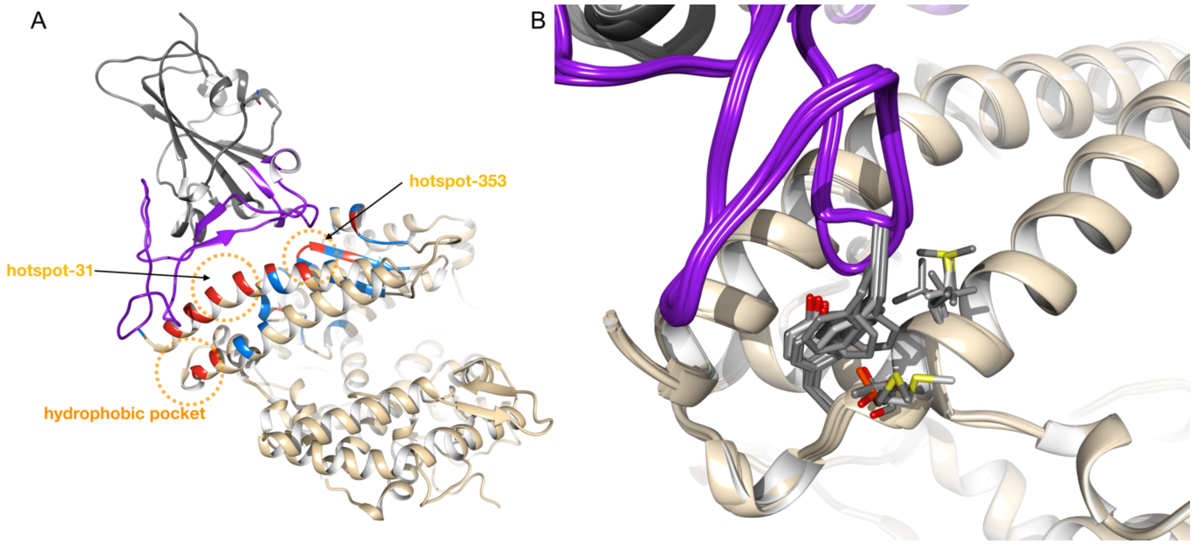


***Supplementary Figure 10: Structural features of the S-protein from SARS-CoV-2 and SARS-CoV bound to ACE2. (a) Three key interaction sites for the SARS-CoV-2 and SARS-CoV S-proteins with ACE2. (b) Variations in hydrophobic pocket residues at the S-protein:ACE2 interface and the range of conformations adopted by RBD residue Phe486 in 6 animal species.***

*Hydrophobic pocket*

The S-protein in SARS-CoV-2 has increased flexibility of the ACE2 interface loop due to a four residue motif (Gly-Val-Glu-Gly) in place of three in SARS-CoV (Pro-Pro-Ala), allowing for a more compact interface and insertion of SARS-CoV-2 RBD's Phe486 into a hydrophobic pocket in human ACE2 comprising Leu79, Met82 and Tyr83[21,22]. We assessed conformations of S-protein Phe486 and these three pocket residues for each of twelve species compared to human. Phe486 binds in a similar conformation with respect to the hydrophobic pocket for all 12 animals analysed (Supplementary Fig. 10b).

*Hotspot-353*

ACE2 Lys353 is highly conserved and a key mediator of the SARS-CoV and SARS-CoV-2 S-protein interfaces[21]. Human Lys353 forms a salt bridge with Asp38 and these residues form H-bonds with the S-protein's RBD (Supplementary Fig. 10a). These interface interactions are significantly disrupted in horseshoe bat which has uncharged residue Asn38 in place of aspartate. This leads to loss of the salt bridge with Lys353 and a change in its conformation that no longer has an H-bond with S-protein Gly496 due to an increase in distance to 4.49Å (Supplementary Fig. 11b).

In cat and dog the physicochemically similar but bulkier glutamate replaces aspartate, leading to loss of both the salt bridge and H-bond interactions at this position. However, we observed a reconfiguration of ACE2 residues permitting alternative H-bonds to the S-protein RBD residues (Supplementary Fig. 11c).

Like the human interface, S-protein is predicted to interact with ACE2 in the primates capuchin, squirrel monkey and marmoset via hotspot-353 as they all conserve Asp38, which can form H-bonds with S-protein Tyr449. (Supplementary Fig. 11d, e & f). Capuchin is predicted to also form the additional H-bond Lys353-Gly496, while squirrel monkey and marmoset have predicted salt bridge Asp38-Lys353 aiding the interface, as in humans, by stabilising the positively charged lysine. However, in koala, a marsupial more distantly related to humans Asp38 is not conserved, though the larger glutamate at the interface is still predicted to form a H-bond with Tyr449.

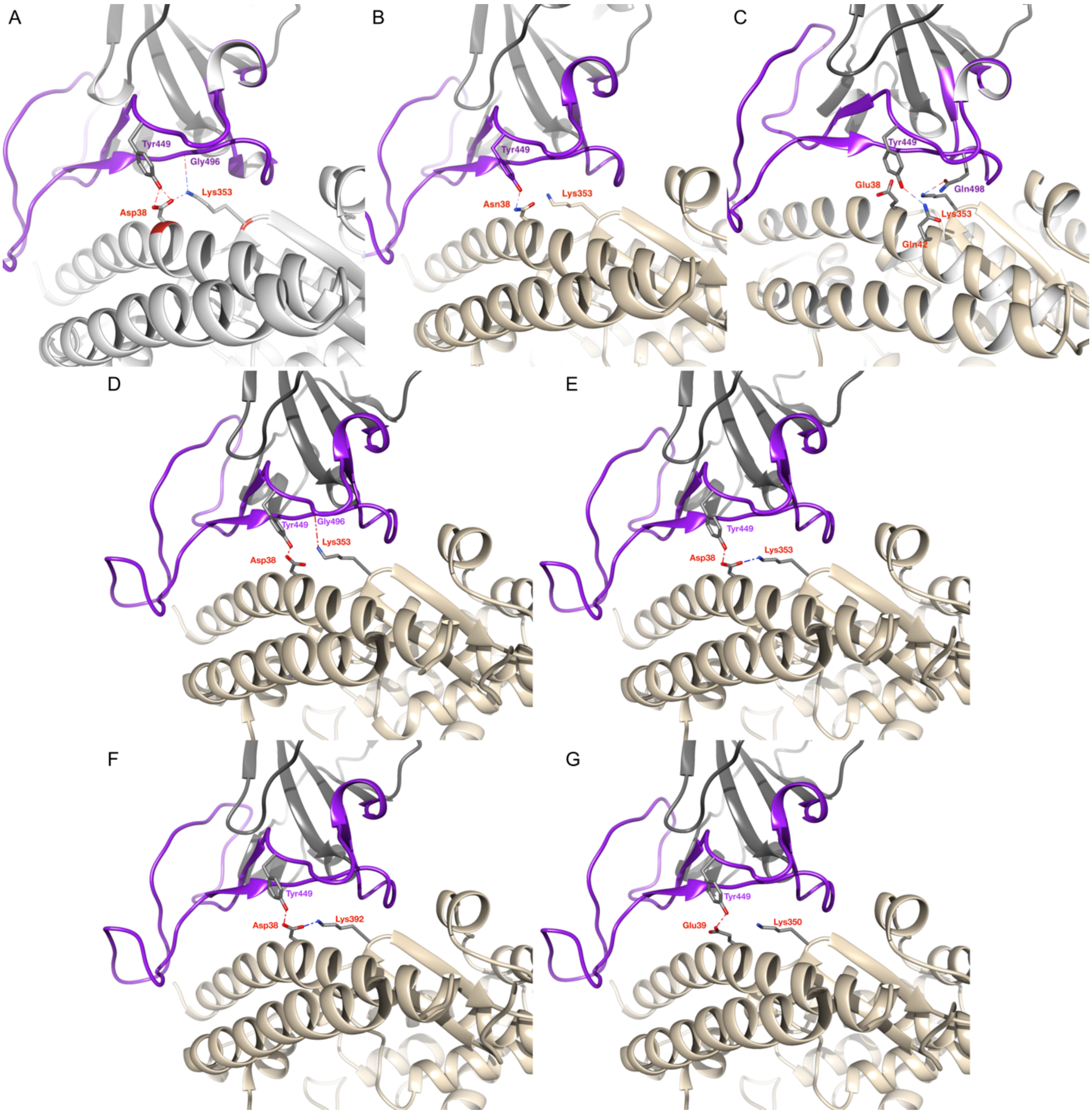


***Supplementary Figure 11: Predicted interactions at the S-protein RBD (purple) interface with ACE2 (tan) around the Lys353 hotspot in human and six other species. (a) In human ACE2 Lys353 forms a salt-bridge with Asp38. ACE2 Lys353 forms H-bonds with S-protein Gly496 and Asp38 with S-protein Tyr449. (b) Horseshoe bat Asn38 only forms H-bond with S-protein Tyr449. (c) In cat and dog bulkier Glu38 no longer forms salt bridge or H-bonds with S-protein, but alternative H-bond interactions are predicted between Lys353 and S-protein Gln498 and Gln42 with Tyr449. (d, e & f) New World Monkeys Capuchin, Squirrel Monkey and Marmoset all conserve Asp38 and H-bond***

***with the S-protein Tyr449 with evidence that either the salt bridge between Asp38 and Lys353 or the H-bond Lys353 to S-protein Gly496 can also occur. (g) In Koala Asp is replaced by Glu39, which can still H-bond with S-protein Tyr449 but predictions did not identify a salt bridge or second H-bond with the S-protein RBD.***

*Hotspot-31*

In this hotspot, ACE2 residue Lys31 is conserved in macaque, cat, cow, dog, sheep, capuchin, squirrel monkey and marmoset with respect to human. The positively charged lysine is replaced with a negatively charged glutamate in guinea pig and camel or aspartate in horseshoe bat, while Koala has uncharged threonine. Glu35 is conserved in 11 of the 12 animals, with only guinea pig having the variant lysine. In human ACE2 Glu35 and Lys31 form a salt bridge which breaks during S-protein binding, then each residue forms H-bonds with S-protein Gln493[21]. Somewhat diverse conformations of ACE2 residues 31, 35 and the S-protein Gln493 were observed in the 12 animal species (Supplementary Fig.12) and H-bonds with S-protein are predicted if the H-bond angle constraint is relaxed by 10 degrees. Finally, the conservation of Glu35 and a charged residue at position 31 in all species except Koala suggest that H-bonding interactions are possible at this hotspot in a wide range of species forming a significant component of ACE2-S-protein binding.

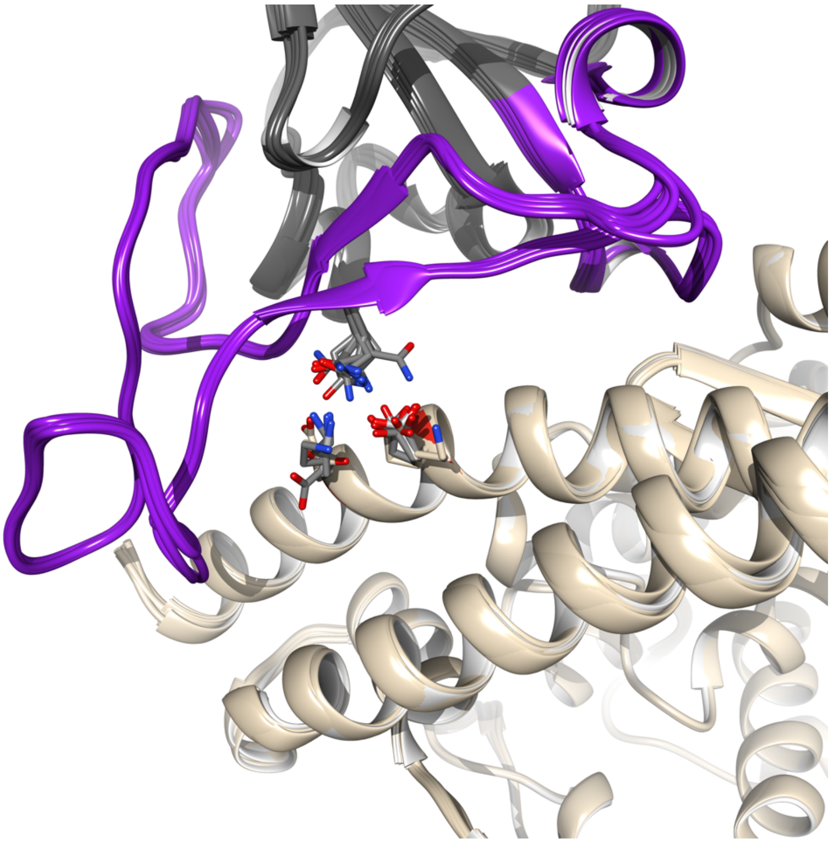

***Supplementary Figure 12: Structural analysis of ACE2 hotspot-31 mutations. Residues Lys31 and Glu35 (in human) and Gln493 from the S-protein RBD, from the human structure is shown with 8 animal species. Glu35 is conserved in all but guinea pig. Variants of human Lys31 are either Asp or Glu and adopt varied conformations.***

*Potential allosteric region formed by non-interface residues*

We identified a cluster of residues in our DCEX set (Supplementary Fig. 13) in close proximity to the well-characterized hotspot residue Lys353 of ACE2. These residues lie in the β3 and β4 strands that are close to the loop containing Lys353. The non-interface DCEX residues that mapped to β3 include Thr347, Ala348, Trp349, Asp350, Gly352, while residues Phe356, Ile358, Leu359, Met360 occur in the β4 strand. In addition, residues Met323, Thr324, Gln325, Ser331, Met332, Leu333, Trp328 and Glu329, map to a helix formed by residues 324-331. Most of these residues are highly conserved with ScoreCons scores > 0.70 and 10 residues (59%) have ScoreCons scores > 0.9. The proximity of the residues to the binding interface suggests a possible allosteric role for these very highly conserved residues.

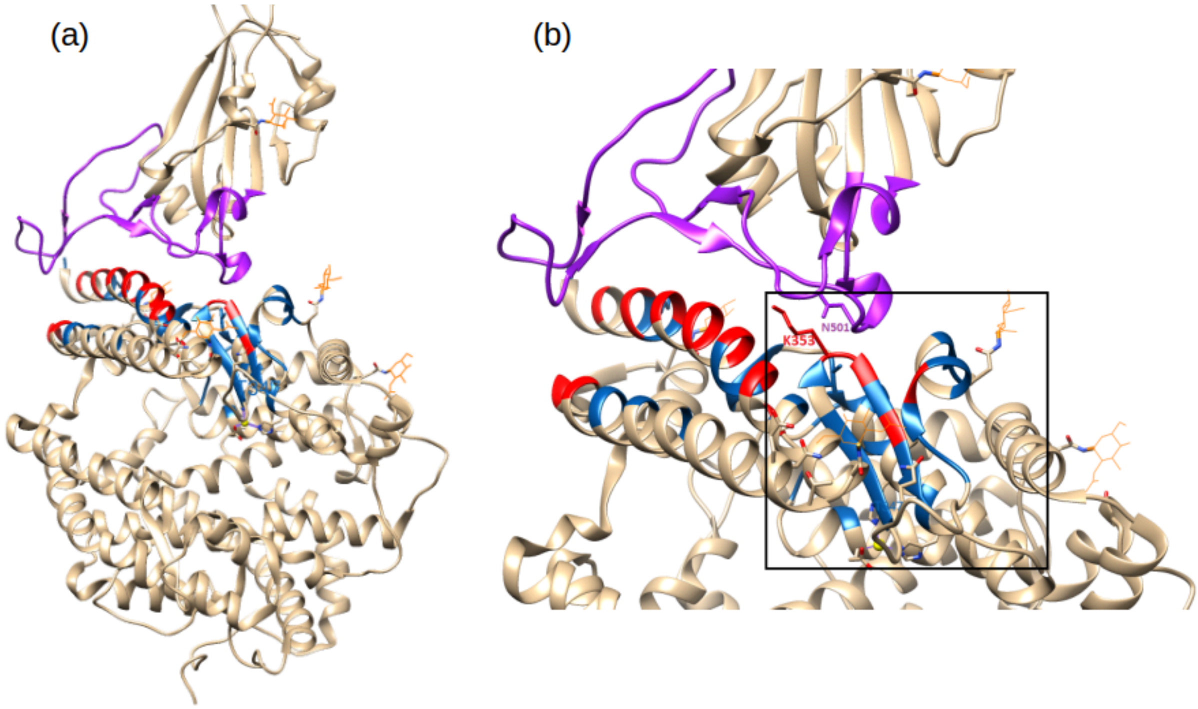


***Supplementary Figure 13: Potential allosteric region, identified in this study: The potential allosteric cluster (shown in blue) formed by non-interface residues is indicated in the box. The cluster is formed near the known hotspot residue Lys353 in ACE2. The cluster is formed by residues (β3: 347-350, 352), (β4: Phe356, Ile358, Leu359 and Met360), which lie close to the loop containing the hotspot residue Lys353. Additionally residues, Met323, Thr324, Gln325, Ser331, Met332, Leu333, Trp328 and Glu329 also form part of the cluster.***

We further tested the allosteric potential of residues in this region using various allosteric prediction methods such as AlloSitePro[24], ENM[25], and PARS[26]. All three methods provide significant evidence for allosteric potential of these residues (Supplementary Fig. 14 and 15). PARS reports significant evidence for the presence of a cavity in this putative allosteric region (residues Thr324, Gly354 and Phe356) with significant flexibility (Mann-Whitney $P \cong 0$).

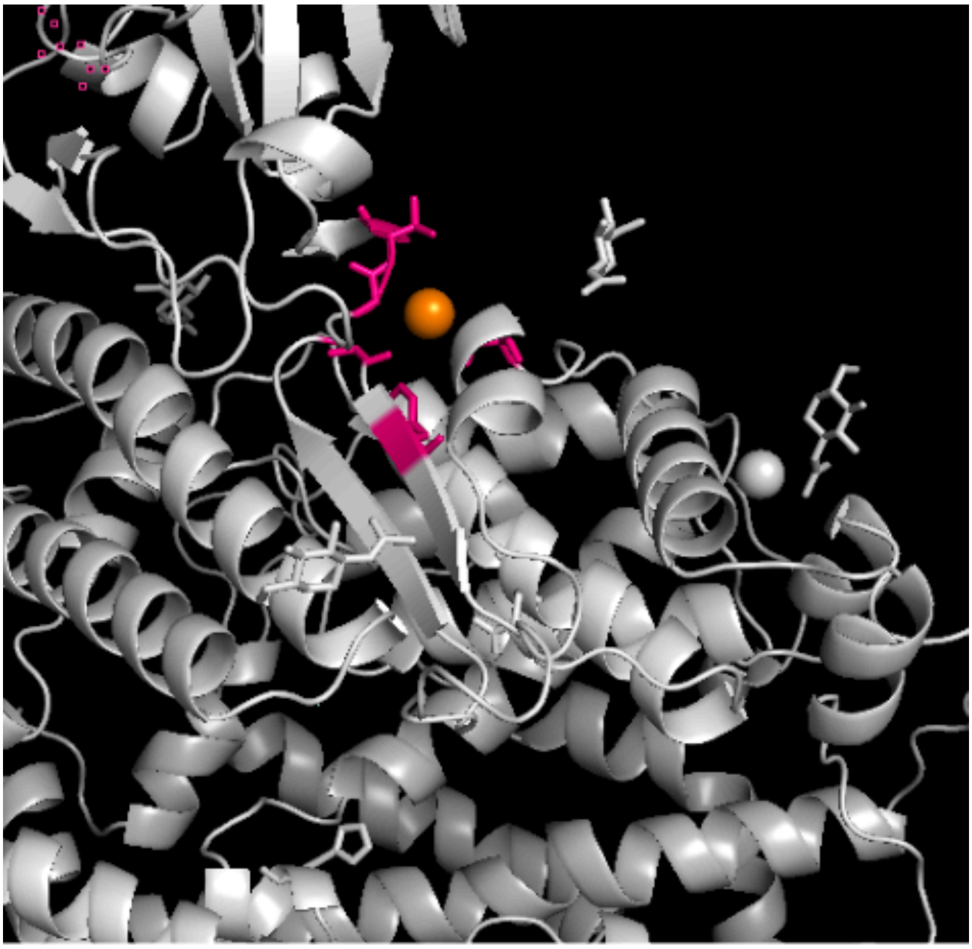

***Supplementary Figure 14: Allosteric sites predicted by PARS method. There is significant evidence for the presence of a cavity in the putative allosteric region (residues Thr324, Gly354, Phe356 shown in pink).It is worth mentioning that PARS provides very high support for this site, as compared to other known binding sites such as active site, binding sites for NAG, CL and Zinc molecules.***

Another method, ENM, reports hinge site positions in the predicted allosteric region, namely Ala348, Phe356 and Met323 (shown below, Supplementary Fig. 15) and also reports Gly395 (Extended non-interface residue), Ile379, Asp382, Asn397, as potential functional sites. Finally AlloSitePro predicts an allosteric site (AlloSite score = 0.507) consisting of non-interface residues, 324-327, 329, 330, 354-356, within the predicted allosteric region. AlloSitePro also identifies a few additional sites such as 320, 321, 380, 383-386, 555 and 558 which are located around this cluster. These methods therefore strongly substantiate this putative allosteric region. In addition, Procko et al.[8] performed deep mutagenesis experiments in ACE2 and highlighted the putative structural roles of certain residues in the secondary shell of the interface including T324 (a DCEX 5Å residue) and Q325 (a DCEX 8Å residue).

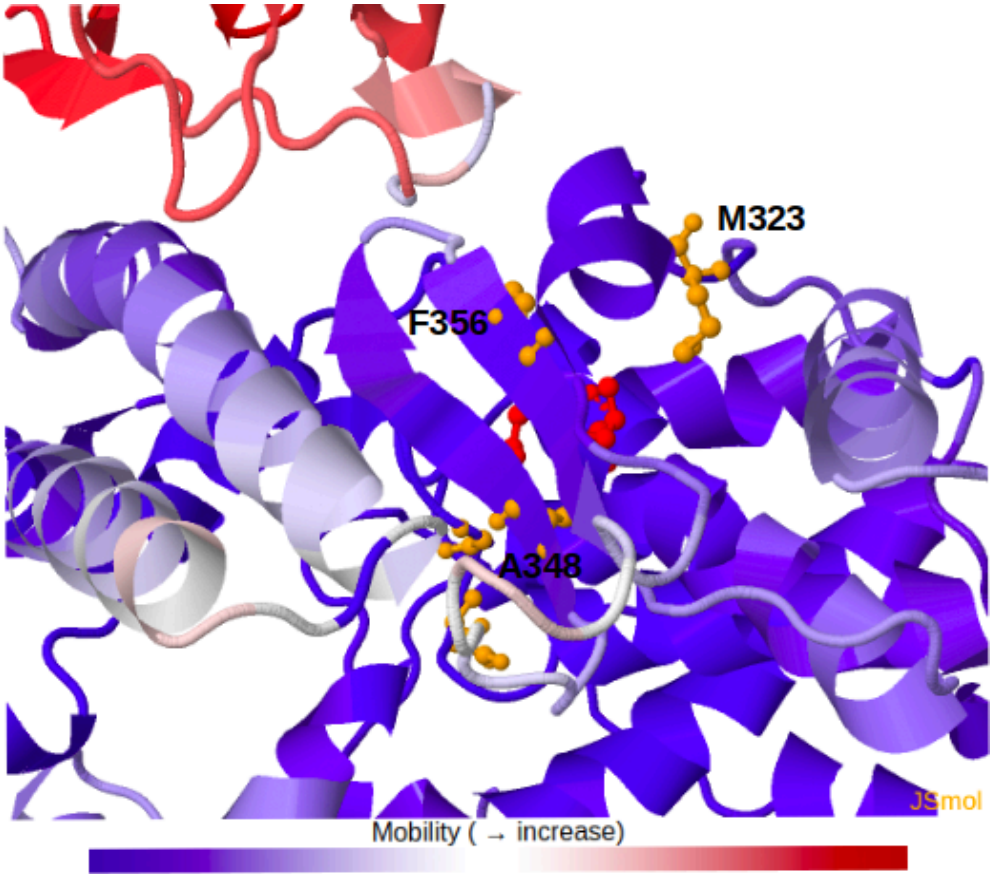


***Supplementary Figure 15: Allosteric sites predicted by DyNOmics ENM method. Potential hinge residues (Ala348, Phe356 and Met323), indicated in red and yellow, are top-ranked residues predicted by the ENM method.***

Although most of these residues are highly conserved across animals, 168 animals have ≥ one mutation and more than 100 animals have ≥ three mutations in this region. However most of these changes are quite conservative. The only significant change is the L359K mutation, which moderately stabilises the complex in many animals.

#### Supplementary Results 5: Changes in energy of the S-protein:ACE2 complex in SARS-CoV-2 and SARS-CoV

Farmed civets, infected by SARS-CoV in the 2002-2004 SARS epidemic[27], are thought to have been intermediate hosts of SARS-CoV(100), and thousands were culled in China to help to control the epidemic[28]. Using the SARS-CoV S-protein sequence, we analysed changes in energy of the S-protein:ACE2 complex for the 215 animals in our study. We used a structure of the SARS-CoV S-protein:ACE2 complex (PDB ID 2AJF) to model structures for the other animal species. Changes in energy of the S-protein:ACE2 complex are highly correlated in SARS-CoV-2 and SARS-CoV (Supplementary Fig. 16 and Supplementary Fig. 17), suggesting that the range of animals susceptible to the virus is likely to be similar for SARS-CoV-2 and SARS-CoV.

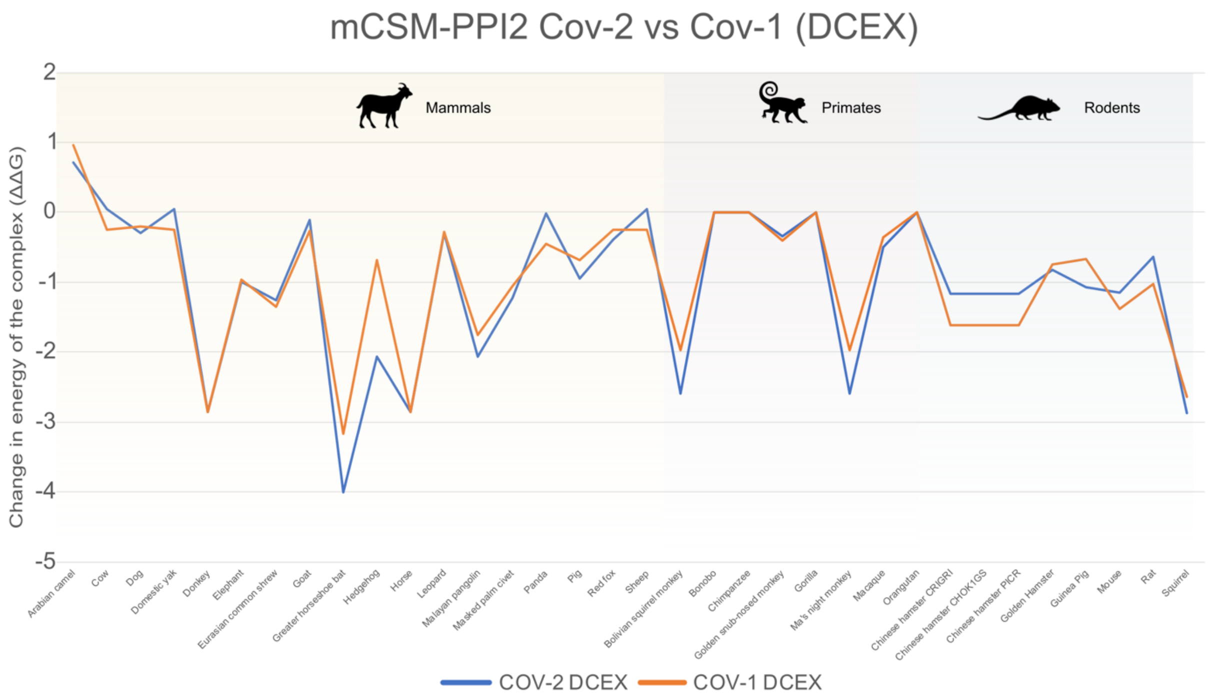


***Supplementary Figure 16: Changes in the energy of the SARS-CoV:ACE2 and SARS-CoV-2:ACE2 complexes across selected animals. The change in energy of the complex (ΔΔG), as measured by protocol (2) - PPI2 over the DCEX residues, is shown on the y-axis.***

It should be noted that the binding of the SARS-CoV-2 S-protein:ACE2 complex is 10-22-fold stronger than for SARS-CoV[5,21,23], however both energies are sufficient for an interaction and clearly did enable infection with human hosts.

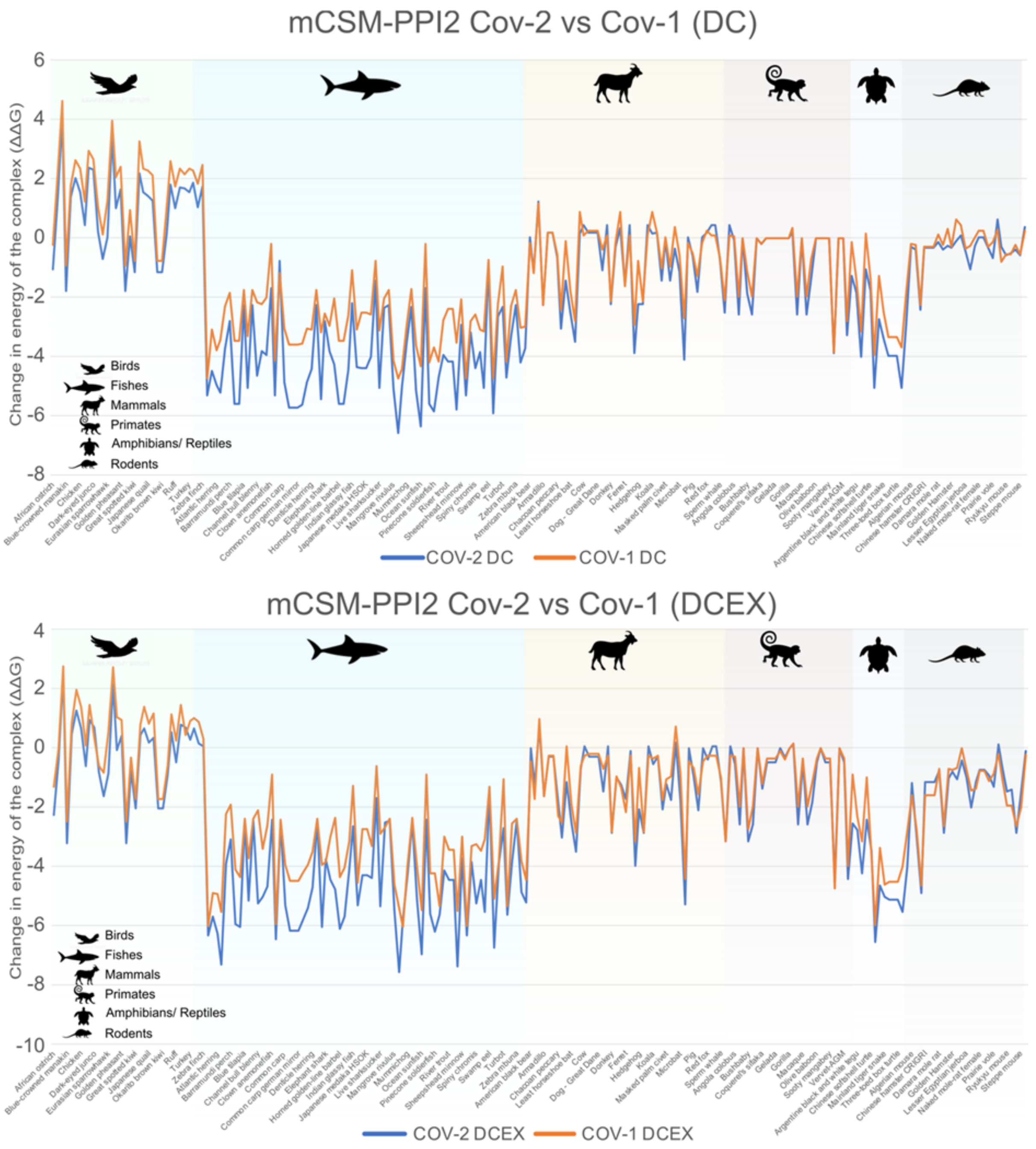


*Supplementary Figure 17: Changes in the energy of the SARS-CoV:ACE2 and SARS-CoV-2:ACE2 complexes across all available organisms. Variation in changes in energy of the S-protein:ACE2 complex across animals is highly correlated for SARS-CoV-2 and SARS-CoV.*

### Supplementary Methods

#### Supplementary Methods 1: Sequence analysis

##### Analyses of sequence similarity between human ACE2 and other vertebrate species

Vertebrate protein sequences were aligned to each other pairwise, and to the human ACE2 sequence, using BLASTp[29].

##### Domain family detection, alignment construction and conservation of residues in ACE2 and TMPRSS2

In order to detect highly conserved residues likely to be implicated in the domain function (i.e. ACE2 binding residues, TMPRSS2 binding residues in the active site) we mapped the domain sequence to the appropriate functional family (FunFam) in the CATH classification[30]. FunFams are clusters of evolutionary related domain sequences, predicted to have highly similar structures and functions. They have been previously used to analyse the impact of genetic variation on protein function in the context of antibiotic resistance phenotypes[31] and cancer progression[32].

The ACE2 (or TMPRSS2) sequences were scanned against the seed FunFam hidden Markov model library for the next release of CATH-Gene3D[33] v4.3 (unpublished data) with the best matches resolved by cath-resolve-hits[34] using a bit score cut-off of 25 and a minimum query coverage of 80%. Each hit was subsequently re-aligned to the matching FunFam using Clustal[35] in Jalview[36].

The conservation scores for residues in the ACE2 and TMPRSS2 domains were obtained using ScoreCons[37]. Information contents of FunFam multiple sequence alignments were calculated as the diversity of positions[37] (DOPs) score. Residues are considered highly conserved for DOPs ≥ 70 and ScoreCons ≥ 0.7.

#### Supplementary Methods 2: Structure analysis

To identify residues involved in the S-protein:ACE2 interface, we extracted information from a range of sources including PDBe[2], PDBsum[3], examined structural evidence (such as from crystallography, cryo-EM and homology modelling) in the literature[4–9], and performed manual inspection of key regions of the human complex (PDB ID 6M0J), identified in previous studies[5,21,22], using UCSF Chimera[38] (Supplementary Fig. 3). Chimera was used for predicting H-bonds, salt bridges and the rendering of struxctural images. Since animal models were built using the slow refinement option in MODELLER[39], side-chain rotamers had been optimised. However, in a few cases we predicted H-bonds at key hotspot residues by exploring alternative rotamers using the Dunbrack rotamer library[40] in Chimera, or by relaxing the allowable H-bond angle constraint. We also examined residues identified by other studies including alanine scanning mutagenesis, deep mutagenesis experiments[5,7,9,41,42], and methods identifying sites under positive selection[43,44].

In addition, in order to identify potential allosteric residues within the ACE2 DCEX set, three different allosteric prediction methods were used: AlloSitePro[24], ENM[25] and PARS[26]. The protein structure PDB ID 6M0J[5] was used as input, which contains information on ligand binding sites for zinc, chlorine, and N-Acetylglucosamine. PARS is a normal mode analysis method which calculates normal modes in the presence and absence of a simulated allosteric modulator. A site is predicted as allosteric if the motions are significantly different. ENM identifies hinge residues as potential functional residues, and relies exclusively on inter-residue contact topology. It uses features of spatial clustering properties and relative solvent accessibility to identify functionally important hinge residues. AlloSitePro uses both pocket-based structure features with normal mode analyses. The performance of AllositePro has been endorsed by recent studies[45].

We also identified DC and DCEX sites under positive selection using codon-based methods, including mixed effect model of evolution[46], available at the Datamonkey Adaptive Evolution web-server[47]. These methods estimate dN/dS ratio for every codon in an alignment. We analysed evidence of positive selection using a codon alignment of all ACE2 orthogue sequences. Potential recombinant sequences were identified using RDP[48] version 5 and were excluded prior to selection pressure analyses.

#### Supplementary Methods 3: Generating 3-dimensional structure models

*ACE2*

Using the ACE2 protein sequence from each species, structural models were generated for the S-protein:ACE2 complex for 247 animals using the FunMod modelling pipeline[33,49], based on MODELLER[39]. Given that the S-protein has the same sequence in PDB ID 6M0J, and ACE2 is conserved in vertebrates, models are expected to be high-quality. Models were refined by MODELLER to optimise the geometry of the complex and the interface. Only high-quality models were used in this analysis, with nDOPE[50] score < -1 and with < 10 DCEX residues missing. This gave a final dataset of 215 animals for further analysis.

*TMPRSS2*

There is currently no solved structure for TMPRSS2. To build a structural model of TMPRSS2, we selected the best TMPRSS2 model template by performing a HMMER[51] search of the human TMPRSS2 sequence (UniProt accession O15393) against the FunFams, identifying four potential structural templates (CATH domain IDs 2f83A05, 5eodA05, 5eokA05 and 5i25A05) within FunFam 2.40.10.10-FF-2 (best E-value = 3e-96). After modelling with FunMod, the structure with the best score (nDOPE = -0.773) was selected, based on PDB ID 5I25.

We identified the relevant residues in TMPRSS2 by extracting the active site and cleavage site residue information available on the human TMPRSS sequence (UniProt accession O15393). After we modelled the human TMPRSS2, two sets of residues were extracted:

1. Set 1: Direct Contact (ASCS) residues comprising the active site (three residues) and the cleavage site (two residues).
2. Set 2: Direct Contact Extended (ASCSEX) residues: This dataset includes the residues from Set 1 and additional residues within 8Å of ASCS residues. We only analysed highly conserved residues (i.e. high ScoreCons ≥ 0.7) from Set 2, which are likely to have a functional role (21 residues).

#### Supplementary Methods 4: Measuring changes in the energy of the S-protein:ACE2 complex in SARS-CoV-2 and SARS-CoV

We used mCSM-PPI1 and mCSM-PPI2 to calculate changes in energy in the SARS-CoV and SARS-CoV-2 complexes following mutations in the DC and DCEX residues. mCSM-PPI1 assigns a graph-based signature vector to each mutation, which is then used within machine learning models to predict the binding energy. The signature vector is based upon atom-distance patterns in the wild-type protein, pharmacophore information and available experimental information. The more recent mCSM-PPI2 is a development of the former with an improved description of the signature vector including, among other features, evolutionary information and energetic terms. For both mCSM-PPI1 and mCSM-PPI2 we used the mCSM server for the simulations.

As an independent validation of our predictions of risk assessment, we also measured the changes in binding energy using the HADDOCK method[20]. HADDOCK is one of the top-performing protein-protein docking servers in the CAPRI competition[52]. We employed HADDOCK (v2.4 web server) to score the complexes. We only performed this for the 41 animals for which we had experimental evidence. We refined the animal 3D models using the default refinement protocol of HADDOCK 2.4[20,53]. Each model was refined through 50 molecular dynamics simulations. These refined models were then clustered using the FCC algorithm[53] with default parameters and scored using the HADDOCK score. The HADDOCK score is a linear combination of van der Waals, electrostatics, and desolvation energy terms. A lower HADDOCK score signifies stronger binding of the S-protein:ACE2 complex.

#### Supplementary Methods 5: Change in residue chemistry for mutations

To measure the degree of chemical change associated with mutations occurring in the key interface residues, we computed the Grantham score[54] for each vertebrate compared to the human sequence. Grantham scores indicate physicochemical similarity between two amino acids calculated using molecular volume, polarity and composition. Grantham scores (range 0-215) are independent of sequence or structural context and we provide them to complement $\Delta\Delta G$ calculations. In general, it would be expected that amino acid substitutions between human and animal species which have a high Grantham score would be more disruptive to interface stability.

For the ACE2-TMPRSS2 comparison we used the same set of organisms as for ACE2, but only those that had at least one TMPRSS2 protein available from ENSEMBL, thus reducing the number of available organisms to 156.

The sums and averages of Grantham scores were obtained for DC and DCEX residues in ACE2. For TMPRSS2 the sum and average Grantham values were obtained for the active site, cleavage site residues (ASCS) and the residues 8Å from the active site (ASCSEX), having ScoreCons ≥ 0.7. Data and plots for the various combinations are available in Supplementary Table 3.

#### Supplementary Methods 6: Phylogeny of SARS-like betacoronaviruses

Genome assemblies were downloaded for a subset of SARS-CoV (n = 10), SARS-like (n = 28) and SARS-CoV-2 (n = 38) viruses from publicly available data on NCBI[55–62] and GISAID[63,64]. The latter were selected to include all high coverage, complete animal associated assemblies together with at least three representatives of human SARS-CoV-2 from each of the seven clades defined by GISAID For a full list of metadata, as compiled by NextStrain[65], including contributing and submitting laboratories, see Supplementary Table 4. All assemblies were downloaded and aligned against the SARS-CoV-2 reference genome Wuhan-Hu-1 (NCBI NC_045512.2) using MAFFT(90). The alignment was manually inspected and the first 130 bp and last 50 bp were masked. A maximum likelihood phylogenetic tree was built on the alignment using RaXML-NG[66] under the GTR+G substitution model with 100 bootstraps to assess branch support. The resulting phylogeny was plotted using ggtree[67] v1.16.6 with branch lengths normalised for ease of visualisation. The spike ORF (21563:25384 nt relative to Wuhan-Hu-1) was extracted and translated to amino acid sequence using Ape[68] v5.3. The alignment of spike protein and receptor binding domain sequences were assessed and visualised using ggtree.
